# Tumour growth: Bayesian parameter calibration of a multiphase porous media model based on *in vitro* observations of Neuroblastoma spheroid growth in a hydrogel microenvironment

**DOI:** 10.1101/2022.09.26.509452

**Authors:** Silvia Hervas-Raluy, Barbara Wirthl, Pedro E. Guerrero, Gil Robalo Rei, Jonas Nitzler, Esther Coronado, Jaime Font De Mora Sainz, Bernhard A. Schrefler, Maria Jose Gomez-Benito, Jose Manuel García-Aznar, Wolfgang A. Wall

## Abstract

To unravel processes that lead to the growth of solid tumours, it is necessary to link knowledge of cancer biology with the physical properties of the tumour and its interaction with the surrounding microenvironment. Our understanding of the underlying mechanisms is however still imprecise. We therefore developed computational physics-based models, which incorporate the interaction of the tumour with its surroundings based on the theory of porous media. However, the experimental validation of such models represents a challenge to its clinical use as a prognostic tool. This study combines a physics-based model with *in vitro* experiments based on microfluidic devices used to mimic a 3D tumour microenvironment. By conducting a global sensitivity analysis, we identify the most influential input parameters and infer their posterior distribution based on Bayesian calibration. The resulting probability density is in agreement with the scattering of the experimental data and thus validates the modelling approach. Using the proposed workflow, we demonstrate that we can indirectly characterise the mechanical properties of neuroblastoma spheroids that cannot feasibly be measured experimentally.

## 1 Introduction

While tumour growth has historically been linked to disruptions at the cellular level alone, in recent years cancer research has recognised the importance of the microenvironment surrounding the tumour [1]. The link between biophysical tumour properties and their surroundings, on the one hand, to signalling pathways in cancer biology, on the other, is therefore crucial to understanding tumour growth and improving cancer treatment.

To mimic the morphological and functional features of *in vivo* tumours within a controlled environment, multicellular tumour spheroids are grown *in vitro* and used as a model [2]. Collagen-based hydrogels are widely used in experiments to create 3D microenvironments. The collagen matrices make it possible to produce matrices with different mechanical properties based on their composition and preparation procedures. There have been several recent attempts to replicate the first stages of tumour formation [2–5]. Microfluidic techniques enable such miniaturisation of tumour growth [6]. Constraining the system to a small scale ensures better control of the environmental conditions. The main advantage of this is that it makes it possible to recreate biological environments that are more realistic than with traditional *in vitro* cell cultures using two-dimensional (2D) culture models [6–8]. Three-dimensional (3D) cell culture models are able to reproduce such features as tumour architecture and metabolism, unlike 2D cultures.

In the present study, we focus on neuroblastoma (NB) cancer. Neuroblastoma is the most common extracranial solid tumour in the paediatric population [9–11]. The median age of diagnosis is 22 months and the primary tumour is usually located in the adrenal glands, inside the abdominal cavity [12]. NB exhibits unique clinical features, including spontaneous regression, but also a high frequency of metastatic disease at the time of diagnosis. This type of tumour is metastatic in 60–70% of cases, with metastasis frequently appearing in bone and bone marrow, liver, lymphatic nodes and, in very young children, even in the eyes or skin [13]. Whereas patients with low-risk diseases have a favourable prognosis of >90% survival, the 5-year survival rate of patients with high-risk diseases is still below 50% [14].

In recent decades, considerable attention has been devoted to computational models that simulate tumour growth, as simulating biological processes facilitates our understanding of the underlying mechanisms of the biological phenomena, particularly in cancer. The hope is that deciphering the biological interactions between cells and the tumour microenvironment provides better tumour therapies and more robust preclinical testing. The most common physics-based models range from macroscopic simulations of volumetric tumour growth [15] to microscopic simulations of the relevant molecular processes [16]. A variety of approaches are used to simulate tumour growth, from discrete or agent-based [17, 18] to continuous models [19]. For studying small-scale phenomena, single-cell models allow a higher degree of spatial resolution, but their computational cost is high. Analysing processes associated with tumour development or growth involves large cell population sizes, and so it is impossible to simulate these processes using discrete models. Hybrid computational models attempt to solve this drawback by coupling continuum models with agent-based models. Phillips et al. [20] proposed a hybrid mathematical approach that characterises vascular changes via an agent-based model while treating nutrients through a continuum model. Similarly, different authors [21, 22] presented a continuum model that included the (pre-existing) vasculature using a one-dimensional approach. We will use a multiphase computational model based on porous media to simulate the growth of the spheroids [23, 24]. The model consists of three phases (namely tumour cells, interstitial fluid and the ECM) plus a number of species, and it is extremely versatile as it is able to simulate a wide range of biological processes.

While experimental research is a powerful tool when it comes to enhancing our understanding of the complex biological processes behind cancer, it can be expensive and time-consuming. And also if time and money would not play a role, even in the best experimental setups both control of conditions as well as measurement possibilities and hence insight are limited. Thus, *in silico* models are presented as a helpful tool for simulating complex experiments, improving our understanding of *in vitro* scenarios, and trialling new ideas in the computational environment prior to *in vitro* experiments [25–27]. Also once the experiments are done, the right computational models—based on first principles—can be extremely useful and can provide additional important insight. But, the synergy between experimental and computational tools even provides powerful opportunities beyond these scenarios. Indeed, the calibration and validation of mechanical models based on experimental data as well as the quantification of the arising uncertainties are integral to scientific activity, especially in a medical context. While computational models have widened their horizons dramatically in recent years, it is crucial to be able to calibrate these models so that relevant patient-specific predictions can be made. In order to do this, the model parameters have to be determined. Some parameters can be measured directly. Since however a direct measurement of most parameters is not possible, they have to be estimated by inverse analysis. There are two general methods of solving the inverse problem: deterministic and probabilistic. While deterministic optimisation techniques yield a point estimate for the best fit, Bayesian methods infer the entire probability distribution, including the uncertainty, which is especially important in a medical setting. Since most models contain a large number of uncertain parameters, which have to be calibrated, it is first necessary to identify the most influential parameters and to distinguish them from non-influential parameters using sensitivity analysis. Subsequent calibration then focuses on the most relevant parameters.

In the field of tumour-growth modelling, several groups have made some advances in this direction. Hawkins-Daarud et al. [28] laid out a Bayesian framework for calibration, validation and uncertainty quantification of tumour-growth models considering synthetic data, and Collis et al. [29] presented the Bayesian calibration of a simple Gompertzian tumour-growth model as a tutorial. More specifically, Lima et al. [30] performed a Bayesian calibration of a stochastic, multiscale agent-based model based on 2D cell cultures of human breast carcinoma cells.

In this paper, we perform a Bayesian calibration of 3D NB spheroid growth based on a multiphase porous medium model and experimental data (see Fig. 1). In Section 2.1, we present the novel 3D NB spheroids experiments performed in microfluidic devices. In Section 2.2, we then summarise the multiphase porous media model of tumour growth. Next, we perform a time-dependent global sensitivity analysis to identify the relative importance of the model parameters, in Section 2.3. This uses the Sobol method [31, 32], which is a variance-based global sensitivity analysis method, combined with a Gaussian process as surrogate model (Section 2.4). We then feed the results of the sensitivity analysis into a Bayesian calibration process based on sequential Monte Carlo (SMC) methods [33] to infer the posterior density of the most influential parameters from the experimental data (Section 2.5). The findings of the study are summarised in Section 3. Finally, in Section 4, we draw conclusions from these results as they relate to potential future research.

**Figure 1:**
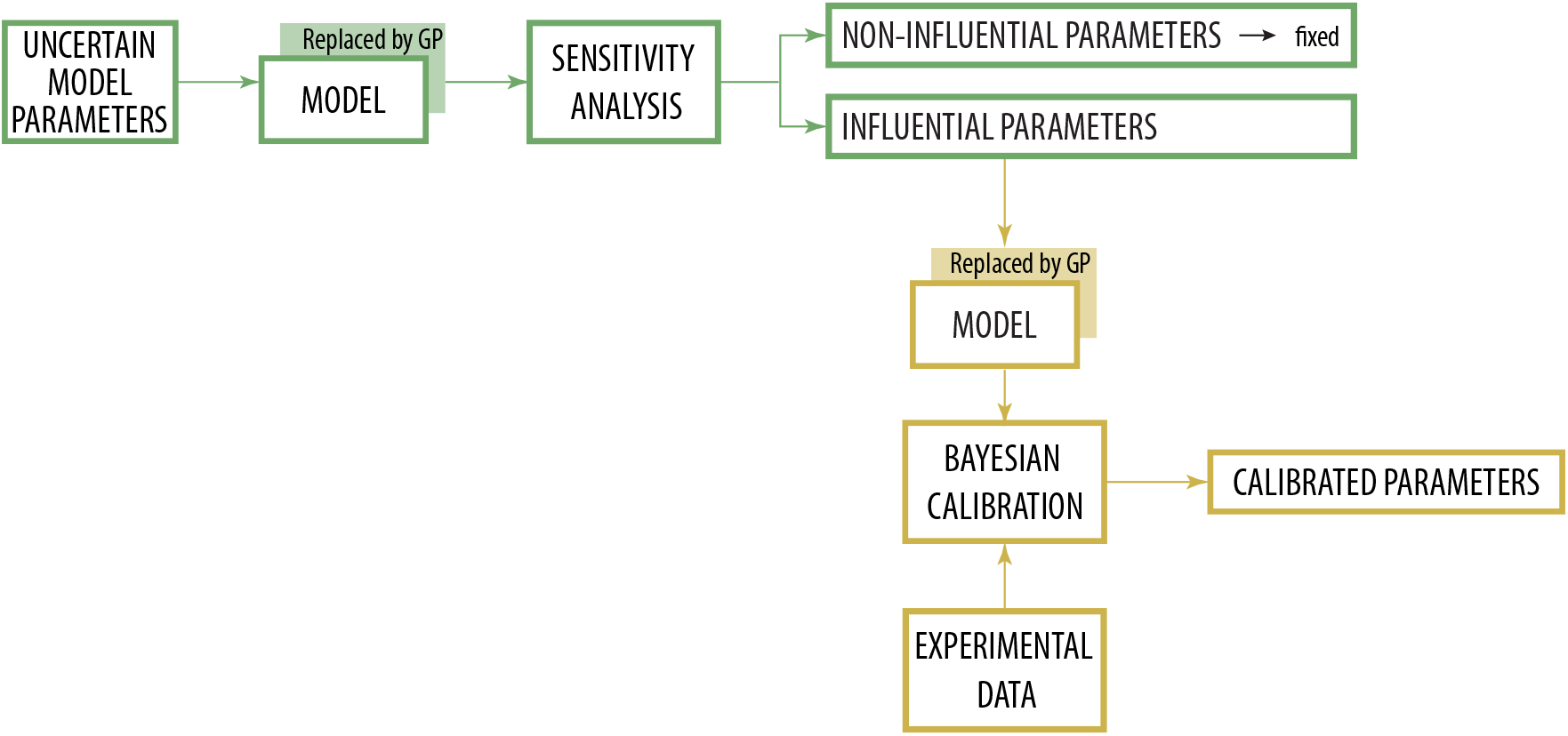
Workflow to calibrate the uncertain model parameters based on experimental data. The uncertain model parameters are first analysed by a global sensitivity analysis, where the model is replaced by a Gaussian process (GP). The sensitivity analysis determines which parameters are non-influential and which have a significant influence on the selected quantity of interest, in this case the spheroid volume. The non-influential parameters are fixed, whereas the influential ones are further calibrated by the Bayesian framework. In this part, the model is again replaced by a Gaussian process, and the influential parameters are calibrated together with the experimental behaviour observed in the microfluidic devices.

## 2 Materials and methods

### 2.1 Experimental setup

The NB tumour spheroids were cultured in 3D microfluidic devices for seven days. We first describe the cell lines used for the experiment and then the experimental setup and image acquisition process. We determined NB spheroid growth from single tumour cells that were homogeneously embedded in 3D-hydrogels.

#### 2.1.1 Cell lines

Neuroblastoma primary culture (PACA cells) was obtained from a patient’s tumour at La Fe University and Polytechnic Hospital (Valencia, Spain), in accordance with the ethics committee of the hospital, reference number 2019-305-1. The patient was diagnosed with poorly-differentiated neuroblastoma harbouring deletion in chromosome 11q, as confirmed by the pathology department of the University of Valencia. The remaining material of the relapsed tumour was used to generate the primary culture. Informed written consent was obtained from legal representatives.

PACA cells were used in 3D-hydrogels in the subsequent experiments. The cells were cultured at 37 °C in a humidified atmosphere of 5% CO_2_. They were routinely grown in Dulbecco’s modified Eagle’s medium (DMEM, Sigma) with high glucose (4.500 mg/mL), L-glutamine and sodium pyruvate and supplemented with 10% heat-inactivated fetal bovine serum (FBS, Gibco) and a 1% antibiotic, antimycotic solution of penicillin, streptomycin, and amphotericin B (Gibco). To enable 3D cell nuclei reconstruction using light-sheet microscopy, the PACA cells were further transfected with a plasmid containing histone 2B fused to GFP (pEGFP-N1 H2B GFP), and clones were selected based on their growth in the presence of G418 (750 μg/mL) and their GFP-expressed fluorescence.

#### 2.1.2 Fabrication of polydimethylsiloxane-based microfluidic devices

The polydimethylsiloxane (PDMS)-based microfluidic devices were produced using the method described by [7]. Briefly, soft lithography was used to develop positive SU8 300-*μ*m relief moulds onto silicon wafers of the desired geometry (National Facility ELECMI ICTS, node *Laboratorio de Microscopías Avanzadas* (LMA) at the University of Zaragoza, Spain). The microdevices were fabricated in PDMS (Sylargd-184, Dow, Midland, TX) at a 10:1 weight ratio of the base and curing agent, respectively. The solution was mixed and poured onto the SU8 master and then degassed to remove any air bubbles. The PDMS was cured for 24 hours at 80 °C, and then the replica-moulded layer was trimmed, perforated with 1 and 4 mm disposable biopsy punches, and sterilized by autoclaving. The PDMS microdevices were activated by plasma treatment and bonded to 35-mm glass-bottom Petri dishes (Ibidi, Gräfelfing, Germany). The microfluidic devices were then coated and incubated for 4 hours at 37 °C with a solution of Poly-D-Lysine (PDL) at 1 mg/mL to enhance adhesion and prevent surface detachment of the collagen hydrogels. After coating, the devices were washed with sterile deionized water and placed in an oven at 80 °C for 24 hours.

The device geometry was described by Plou et al. [2] and is illustrated in Fig. 2. It has a central chamber with a height of approximately 300 μm, a width of 1300 μm, and contains an array of trapezoidal posts to cage the collagen hydrogel solution with the embedded cells. Concerning the central chamber (which was filled with hydrogel), our devices also had two side media channels (each with two media reservoirs) for hydration and medium replacement and to assure the transport of nutrients and other chemical factors throughout the hydrogel.

**Figure 2:**
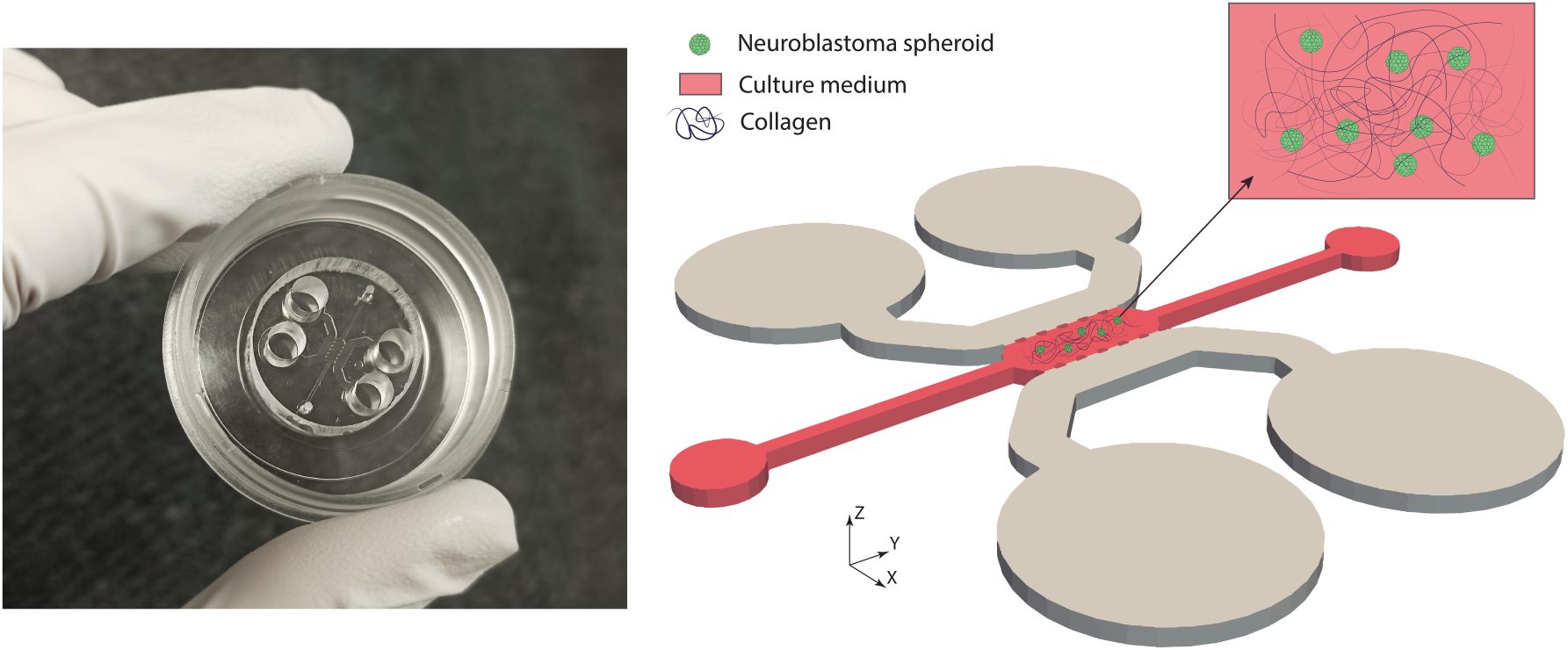
Microfluidic devices used for the analysis of 3D NB spheroids. The image on the left shows the microfluidic device bonded to a 35 mm glass plate. On the right is a 3D visualisation of the microfluidic chip system. A blow-up of the central channel is shown in the top-right. Collagen-I hydrogel is loaded through the central cage (red). The setup includes two side media channels (grey) to supply the culture media and ensure hydration and transport of nutrients and other chemical factors throughout the hydrogel. The height of the culture chamber is 290±20 μm.

#### 2.1.3 Collagen gel solution preparation and cell seeding

We followed the protocol described by Shin et al. [7] for the collagen hydrogel preparation. Collagen Type-I (BD bioscience) was buffered to a final concentration of 6 mg/mL at pH 7.4 with 250,000 cells/mL using 10X phosphate buffered saline (PBS) (Thermo Fisher) with phenol red, 0.5N NaOH solution and DMEM 10%FBS (Gibco). After the central chamber was filled with gel solution, it was left to polymerize for 20 min in humid chambers in a CO_2_ incubator at 37 °C. After polymerization, the gels were hydrated with complete media and stored in the incubator. Media were aspirated from all reservoirs of the device and replaced every 48 hours. Cells were cultured up to day 7.

#### 2.1.4 Brightfield image acquisition and analysis

Cell spheroid formation and growth were visualized and recorded every 24 hours using a Leica DM IL LED inverted microscope in a Basler acA1920-155um camera. Phase-contrast time-lapse images were taken using a 4X Leica objective lens. Images were converted to hyperstacks and aligned with a template matching ImageJ plug-in (National Institutes of Health, USA). The aligned images were then analyzed with a semi-automatic in-house MATLAB code [34] capable of recognising spheroids and calculating their area (μm^2^) and solidity over time. Figures were designed with Prism7-GraphPad (GraphPad Software Inc., San Diego, CA, USA).

#### 2.1.5 Immunofluorescence staining and imaging

For confocal imaging, neuroblastoma spheroids were fixed and immunostained on day 7. Devices were fixed for 15 min at room temperature with 4% paraformaldehyde in PBS. The microfluidic devices were then washed three times with PBS and permeabilized for 20 min with 0.1% Triton X-100 (Calbiochem) diluted in PBS. Devices were washed three more times with PBS and blocked in PBS 5% bovine serum albumin (BSA, VWR) overnight at 4 °C. The samples were then incubated in darkness for a minimum of two hours with phalloidin-tetramethylrhodamine B isothiocyanate (Santa Cruz) and DAPI (Invitrogen). Finally, the devices were washed three more times and imaged. Confocal images were captured at the Microscopy and Imaging Core Facility, *Instituto Aragonés de Ciencias de la Salud* (IACS) with a Zeiss LSM 880 confocal microscope using a 25x oil immersion objective lens. Spheroid projections were obtained from a collection of multiple focal planes (Z-stack). For *in vivo* nuclei visualization, NB spheroids with PACA-GFP (transfected with a plasmid containing histone 2B fused to GFP) were imaged with the Lattice Lightsheet 7 microscope (Zeiss) in a manner that is compatible with long term fluorescent time-lapse imaging. Images were acquired with a 40x water immersion objective lens, processed, and 3D projected using Zen 3.5 Blue software (Zeiss).

### 2.2 Porous multiphase tumour-growth model

A computational model able to reproduce the *in vitro* experiments is now needed. We propose to use a multiphase porous media model of the type that has been widely used to model tumour growth [21, 35, 36]. From a computational modelling perspective, the above experiments can be described as follows: the ECM constitutes a solid porous scaffold with the tumour cells and the culture medium filling the pore space. Together, they form the different phases of a porous medium (see Fig. 3, left). We also include several species, namely oxygen (as the nutrient which drives the growth of the tumour) and necrotic tumour cells (which are formed when deprived of nutrients). At the microscopic scale, all of these components are clearly discernible and distinguishable. We now consider the porous medium at the macroscopic scale, where averaging or up-scaling the microscopic balance equations leads to a continuous description of the porous medium. The solid (in this case the ECM) and the fluids (tumour cells and interstitial fluid) are modelled as overlapping continua, and thus the actual interfaces between the components are not resolved explicitly. At the macroscopic scale, the different phases are described by their volume fractions *ε^α^* at a specific point (see Fig. 3, right), where *α* ∈ {*s*, *t, l*}. In the following, the ECM is denoted by a superscript *s*, tumour cells by *t* and the culture medium by *l*^1^. Note that we model the growing tumour spheroid itself as a viscous fluid, as in the previous tumour models [23]. When averaging the equations, we maintain a strong connection between the microscopic and the macroscopic descriptions by employing the thermodynamically constrained averaging theory (TCAT) [42]. This has the additional advantage that all quantities at the macroscale have a clear relation to their counterparts at the microscale and therefore lend themselves to precise physical interpretation [23].

**Figure 3:**
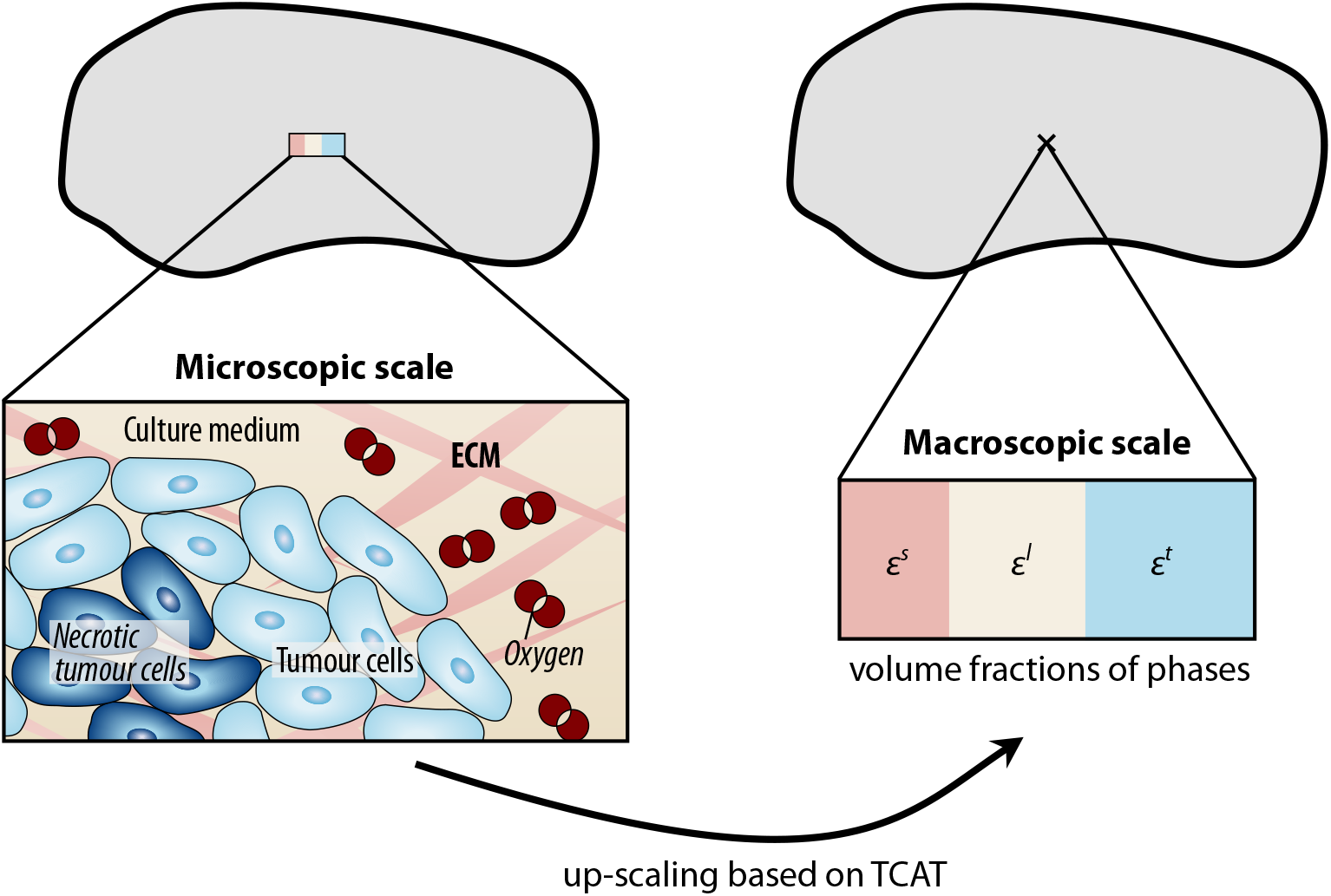
Microscopic to macroscopic up-scaling. At the microscopic scale, all components (ECM, cells, culture medium and oxygen) are resolved. Up-scaling results in a continuous description at the macroscopic scale, in which the phases are described by their volume fractions *ε^α^*. Employing the thermodynamically constrained averaging theory (TCAT) ensures a firm connection between the two scales.

The resulting tumour-growth model has previously been presented in various forms [21, 23, 24, 35–41]. To replicate the spheroid growth experiments, the model is reduced to its two-phase model form, comprising the ECM as the solid phase with two fluid phases flowing in its pores, tumour cells and interstitial fluid. In addition, all phases can transport species. Oxygen, as a nutrient, is assumed to be transported by the interstitial fluid (IF). A sufficiently high oxygen supply leads to tumour growth. In contrast, tumour cells become necrotic when exposed to low nutrient concentrations or excessive mechanical pressure. To take this into account, tumour cells are divided into living and necrotic tumour cells, the latter being modelled as a species.

In the following, we present an overview of the components of the tumour-growth model and its essential input parameters. All model equations are listed in Appendix A.

#### Solid phase: ECM

The ECM is the solid phase of the porous system. Experimental characterisation of the mechanical properties of the hydrogels used in the experiments revealed that hydrogels are usually viscoelastic [43–45], and so we employ a visco-hyperelastic constitutive law. The hyperelastic part is described by a Neo-Hookean material model with the initial Young’s modulus *E* and a Poisson’s ratio *v*, while the viscous part is added in the form of a Maxwell model based on [46] with a relaxation time *τ*. The governing equation of the solid phase is the classical balance of momentum (see Appendix A for further details).

The ECM contains pores (voids) which are filled by the fluid phases (tumour cells and culture medium). The ratio of the ECM pore volume to the total volume is characterised by the porosity *ε*. We assume that the two fluid phases completely fill the pore volume, and hence the porosity corresponds to the sum of the volume fractions of tumour cells *ε^t^* and of the culture medium *ε^l^*:

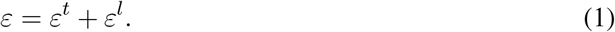

The saturation of the fluid phases is given by

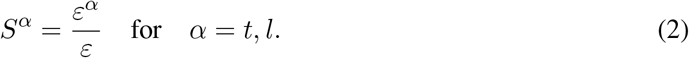

The initial porosity and initial saturation of tumour cells are denoted by *ε*_0_ and 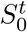, respectively. Since the ECM is deformable, the porosity, volume fractions of the fluid phases, and saturations change as the tumour grows.

#### Fluid phases: culture medium and tumour cells (TC)

We consider two fluid phases that fill the pore space of the ECM: culture medium and tumour cells. The culture medium is modelled as a fluid with a low viscosity *μ^l^*, similar to that of water. The tumour cell phase has a considerably higher viscosity *μ^t^*. The two fluid phases are further characterised by their density *ρ^α^*. The governing equations of the fluid phases are mass balance equations based on the pressures *p^l^* and *p^t^* of the culture medium and the tumour cell phase, respectively (see Appendix A). As the two fluid phases share the pore space of the ECM, they interact with each other and with the ECM. These interactions are influenced by permeability and interfacial tension.

Permeability describes how easily a fluid can flow through the solid scaffold. We define the permeability tensor as 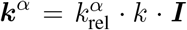 with the scalar intrinsic ECM permeability *k*. The relative permeability 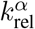 includes interaction forces between the fluid phases and is given by

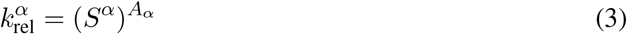

with a model coefficient *A_α_*, as proposed by [38]. The relative permeability is a function of the corresponding saturation, since it depends on the configuration of the two phases within the pore space [47].

Although they are adjacent and share one pore space, the two fluid phases have different pressures due to the interfacial tension *σ_tl_* [35]. This pressure difference *p^tl^* is given by

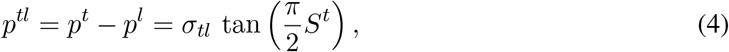

which provides the link back to the fluid saturations. These relationships are similar to those of Parker and Lenhard [48, 49] commonly used for geophysical three-phase systems. A high interfacial tension causes higher infiltration of one fluid phase into the other and hence a less compact tumour [38].

#### Species: oxygen and necrotic tumour cells

The fluid phases can transport further subcomponents. We include oxygen as a nutrient and necrotic tumour cells. The lack of nutrients causes living tumour cells to become necrotic, and these necrotic tumour cells are a subcomponent of the tumour cell phase. The mass fraction of necrotic tumour cells is denoted by 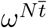. Similarly, nutrients are modelled as a species in the culture medium. The nutrient concentration drives the growth of the tumour. To begin with, we only include oxygen as a nutrient, with the mass fraction 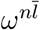. We describe the diffusion of oxygen in the culture medium by 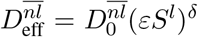, with the diffusion coefficient in the medium 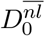 and a constant *δ*. The effective diffusion coefficient 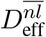 of oxygen has a nonlinear dependence on the volume fraction of IF, as it is also related to the connectivity grade of the extracellular spaces and tortuosity of the porous network. The parameter *δ* was calibrated experimentally by Sciumè et al. [35]. The governing equations used for the species are reaction-diffusion-advection equations (see Appendix A).

#### Mass transfer terms governing tumour growth

To bring everything together, we are left with the question of what actually drives the growth of the tumour in the multiphase porous media model: as long as the tumour cells are supplied with enough oxygen (above the critical threshold 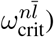, the tumour phase grows, as described by the mass transfer term from the culture medium to the tumour phase

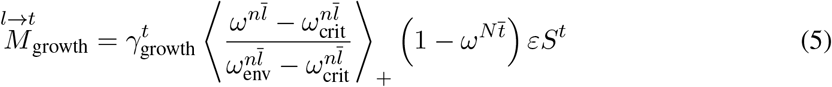

where 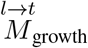 represents the inter-phase exchange of mass between the phases *l* and *t* (representing the mass of IF which becomes tumour due to cell growth) and 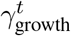 denotes the growth coefficient [35]. The parameter 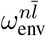 is the mass fraction of oxygen available in the environment, and 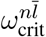 is the critical nutrient threshold below which cells starve and become necrotic. The Macaulay brackets 〈·〉_+_ indicate the positive value of the argument if the argument is positive but zero if it is not.

Moreover, living tumour cells consume oxygen, as described by

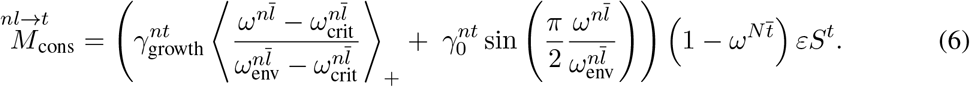

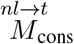 represents the inter-phase exchange of the mass of oxygen present in the IF which becomes tumour due to cell consumption. The first addend describes the consumption of oxygen during tumour growth proportional to the coefficient 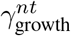, and the second addend accounts for the normal metabolism of tumour cells proportional to the coefficient 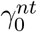.

Finally, tumour cells become necrotic when not supplied with enough oxygen. This increase in the necrotic fraction of tumour cells is included as

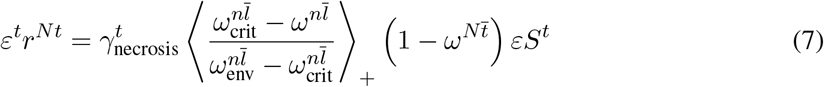

where *ε^t^r^Nt^* is the death rate of tumour cells and 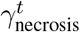 is the necrosis coefficient.

#### Geometry and initial and boundary conditions

The model aims to reproduce the experimental assays described in Section 2.1. The spheroids are assumed to be spherical, with an initial mean radius of 10 μm. Due to the axial symmetry of the growing spheroids, only a segment of a spherical 3D geometry is computationally resolved. The total spherical computational domain has a radius of 100 μm, and we only consider a segment of 0.16 rad × 0.16 rad. Tumour cells initially occupy the inner part Ω^*t*^ with a radius of 10 μm. This simulates the initial cell seed in the microfluidic device. At the initial time, the tumour saturation is set to *S^t^* = 0.875, and zero in the rest of the domain. In addition, we assume that there are no necrotic cells at the initial stage. We assume a steady nutrient supply since the culture medium in the experiment is changed every two days in order to ensure a steady nutrient concentration and a physiological pH level. Therefore, the mass fraction of nutrients in the medium is set as constant and equal to 4.2 · 10^−6^ [41]. It is applied as a Dirichlet boundary condition at the outer surface Γ of the domain. The domain is fixed at the inner surface with a Dirichlet boundary condition but can deform in the radial direction. Fig. 4 summarises the initial and boundary conditions of the numerical analysis.

**Figure 4:**
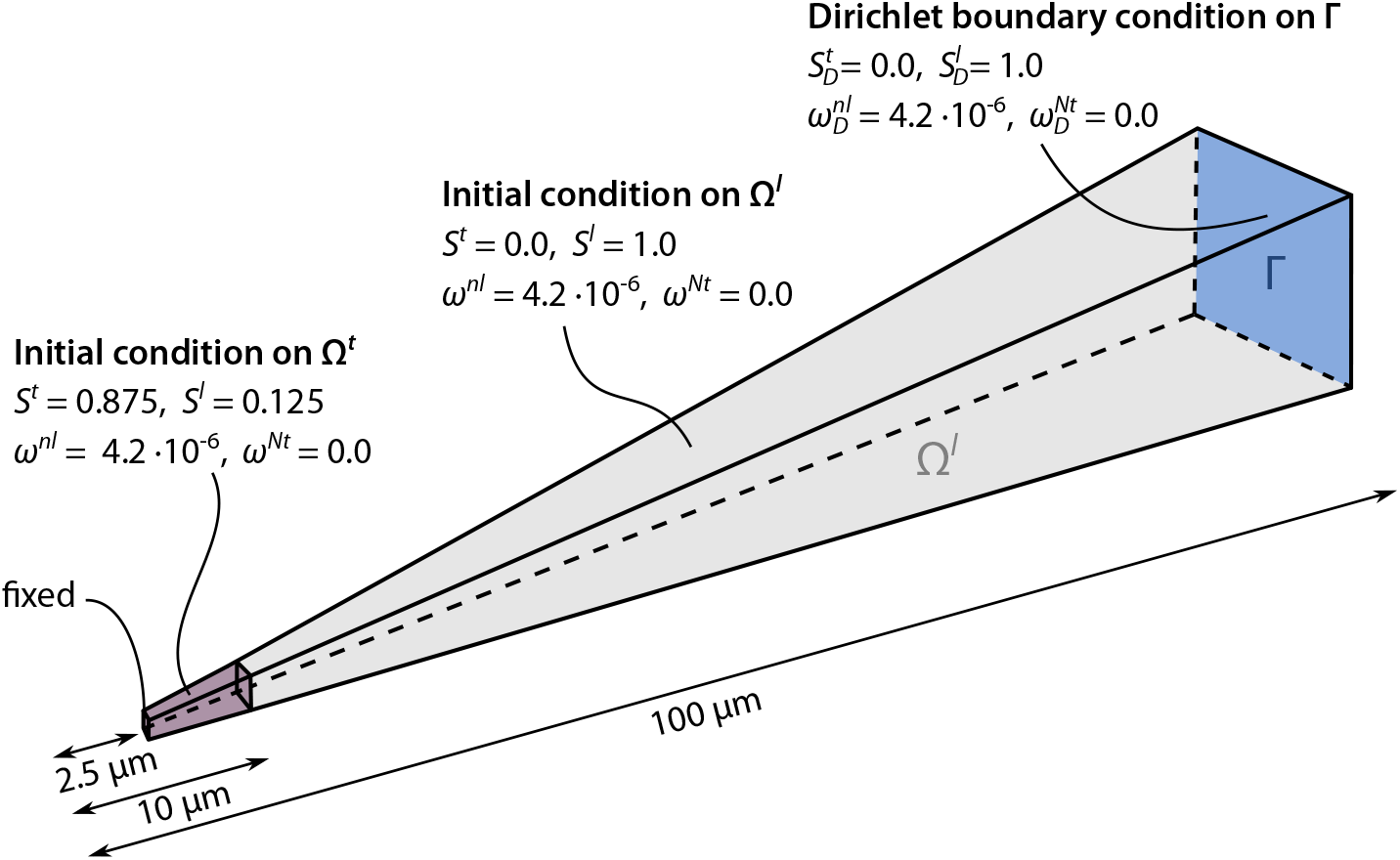
Initial and boundary conditions in the simulated domain. The tumour domain Ω*^t^* is marked in purple, and the domain Ω*^l^* filled with culture medium is marked in grey. Dirichlet boundary conditions for the fluid phases and species are applied on the outer surface marked in blue.

The geometry is discretised with 250 3D trilinear hexahedral elements in the radial direction. The time step is Δ*t* = 450 s. In total 1344 time steps are simulated, and thus the simulation describes the growth of a spheroid over the course of seven days.

### 2.3 Global sensitivity analysis (GSA)

Our tumour-growth model, as described in Section 2.2, is based on physical laws, unlike many other tumour-growth models, which are purely data-driven. Such a physics-based approach allows the model output to be predictive, even under unobserved conditions, but *per se* comes at the cost of a large number of uncertain input parameters. It is therefore crucial to be able to distinguish influential parameters from non-influential ones. Sensitivity analysis does this by quantifying the influence of each uncertain input parameter on the variation of the model output [50, 51], thereby guiding the subsequent calibration process. To perform a rigorous global sensitivity analysis, we employ the Sobol method [31].

#### 2.3.1 The Sobol method

The Sobol method [31, 52] is a variance-based global approach, which decomposes the output variance into portions attributed to the specific input parameters. Being a global approach, the Sobol method explores the entire input space of the uncertain input parameters as opposed to local sensitivity measures, which only perturb a single input parameter at a time around a fixed base point [51].

Our tumour-growth model (hereafter referred to as *f*) includes a number of uncertain input parameters *θ_i_*, which we further detail below. The uncertain input parameters *θ_i_* are assumed to be independent random variables described by their probability density function (PDF) *p*(*θ_i_*) and summarised in the random vector ***θ***, such that 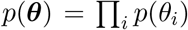. By ***θ***_~*i*_, we denote the random vector of all components except *θ_i_*. Because of the randomness in the input parameters, the output *Y* of the model *f* is also a random variable: *Y* = *f*(***θ***). We use the expected value of the function *f* of ***θ*** denoted by

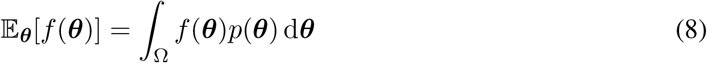

and the variance (second moment) denoted by

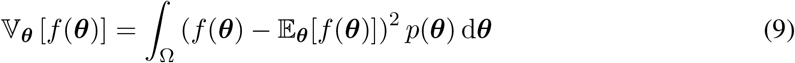

where Ω is the input parameter space.

The first-order Sobol index 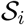 then is defined as^2^

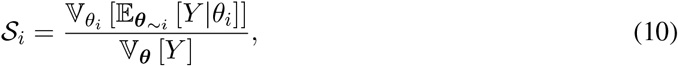

and describes the extent to which the variance of the output 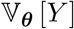 would be reduced if the true value of the input parameter *θ_i_* was known (with 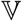 denoting the variance operator) [53]. Note that the expectation 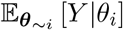 in the numerator is taken over all possible values of ***θ***_~*i*_ (denoted by the subscript) while keeping parameter *θ_i_* fixed. In contrast, the variance 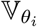 in the numerator is taken over all possible values of *θ_i_*. A parameter *θ_i_* with a high first-order Sobol index 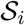 is considered in the subsequent calibration, since determining its true value efficiently reduces the overall uncertainty of the model output, in this case, the tumour volume [50]. At the same time, a small first-order index is not a sufficient basis from which to conclude that the parameter *θ_i_* is non-influential, because the parameter may be involved in interactions with other parameters.

We therefore additionally consider the total-order Sobol index

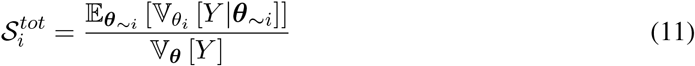

which describes the total contribution of the parameter *θ_i_*, including the first-order effect plus any higher-order effects that arise from interactions. If 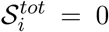 (or, in practice, sufficiently small), the parameter *θ_i_* can be considered non-influential, and hence it can be fixed anywhere in its input range without affecting the output variance [50]. The difference 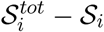 between the first- and the total-order Sobol index quantifies the interactions between the input parameter *θ_i_* and all other input parameters.

To estimate the Sobol indices, we need to compute the conditional expectations and variances given in Eqs. (10) and (11), which means evaluating the multidimensional integrals in the space of the input parameters. Monte Carlo integration is employed for this purpose, as the integration error is not dependent on the dimension of the integrand, unlike grid-based integration schemes, such as Gaussian quadrature [54]. We further use the estimators proposed by [55, 56]. However, this estimation involves evaluating the tumour-growth model *f* for all Monte Carlo samples. Considering the large number of Monte Carlo samples needed and the computational cost of the model *f*, the degree of resources required makes this option unfeasible. We therefore substitute the simulations of the forward model *f* with realisations of a trained Gaussian process metamodel *f*_GP_ (see Section 2.4). The estimation of the Sobol indices includes two main sources of uncertainty: one related to the Monte Carlo integration, and one related to the metamodel approximation. We quantify those uncertainties based on [57, 58].

#### 2.3.2 Setup of the GSA

The quantity of interest in the spheroid growth analysis is the volume *V^t^* of the tumour spheroid over time (see supplementary Eq. (A11)), and the objective of the sensitivity analysis is to separate the (most) influential uncertain input parameters from the non-influential ones. To this end, the input parameters are grouped according to whether they are fixed or uncertain.

The fixed parameters are listed in Table 1. We assume that the values of those parameters are known *a priori*, e.g., from previous studies. The initial tumour radius *r*_0_ is obtained from the experimental images. The densities of the fluid phases (*ρ^t^* and *ρ^l^*) and of the ECM (*ρ^s^*) are assumed to be equal to the density of water. In the experiments, nutrients are supplied every day to ensure that they are available in sufficient concentration throughout the assay. In the computational model, we include oxygen as the only nutrient, and assume that the parameters related to oxygen (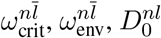 and *δ*) are known.

**Table 1:**
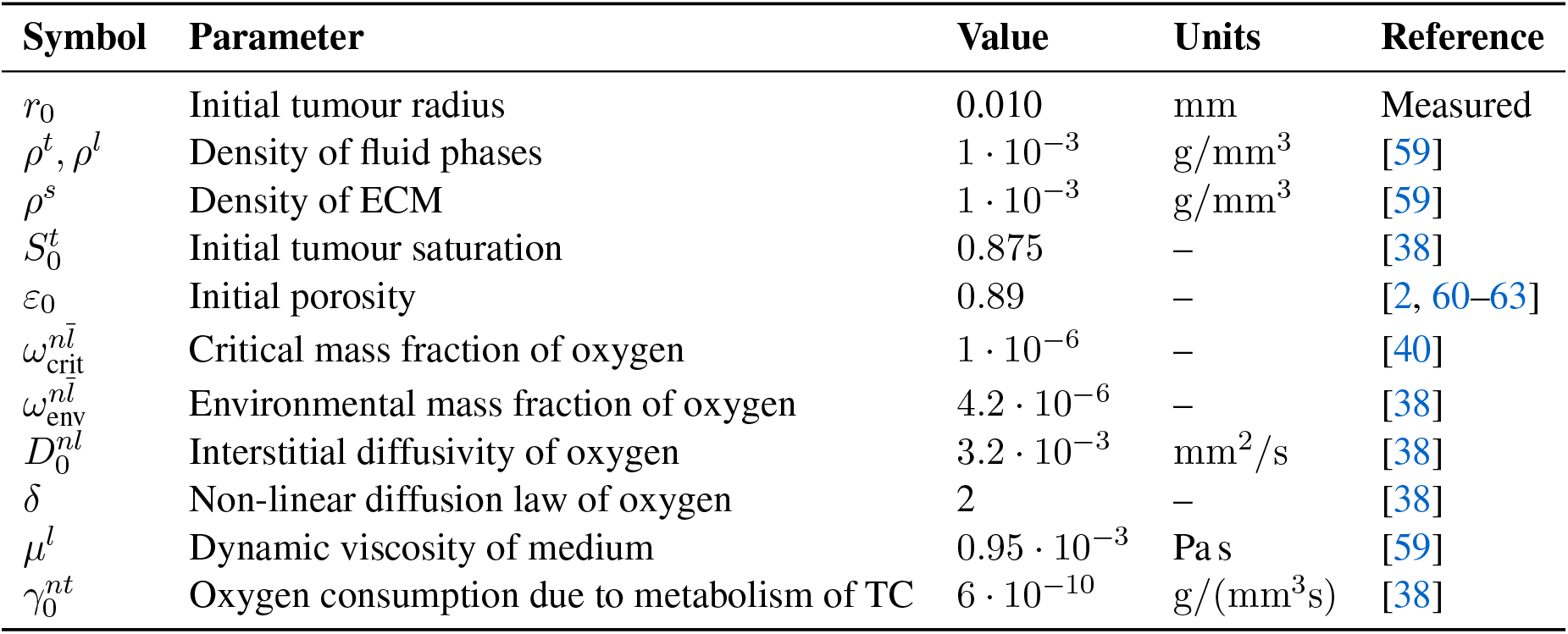
Fixed parameters of the model.

Table 2 lists the eleven uncertain parameters whose influence will be analysed in the GSA. We assume a uniform distribution for all parameters, with 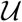 denoting the continuous uniform distribution over the interval [*a*, *b*]. We define the support for the distributions of these parameters based on values obtained from the literature. The ranges of the initial porosity *ε*_0_ and of the properties of the ECM (intrinsic permeability *k*, shear modulus *G*, Poisson’s ratio *ν* and dynamic viscosity *μ^s^*) are based on experimental results, in which the role of the ECM was investigated. The parameters *A_t_* and *A_l_* impact on the isotropic permeability of the ECM with respect to the fluid phase *α* ∈ {*t, l*}. The dynamic viscosity of the tumour cell phase *μ^t^* has been measured in numerous studies, producing values that vary over several orders of magnitude. Further parameters, such as the initial tumour saturation 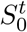, the interfacial tension between the tumour cells and the medium *σ_tl_*, and the coefficients related to growth, necrosis or nutrient consumption (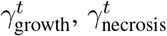 and 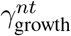) have been investigated in previous publications.

**Table 2:** Probability distributions of the uncertain input parameters.

| Parameter |  | Distribution | Units | Reference |
| --- | --- | --- | --- | --- |
| Interfacial tension | $\sigma_{tl}$ | $\sim \mathcal{U}[2, 50]$ | mN/m | [38, 64] |
| Dynamic viscosity of TC | $\mu^t$ | $\sim \mathcal{U}[10, 1000]$ | Pa s | [24, 65–71] |
| Relative permeability exponent for TC | $A_t$ | $\sim \mathcal{U}[1, 10]$ | – | [41] |
| Relative permeability exponent for medium | $A_l$ | $\sim \mathcal{U}[1, 7]$ | – | [41] |
| Tumour-growth coefficient | $\gamma_{\text{growth}}^t$ | $\sim \mathcal{U}[0.1, 2.0] \cdot 10^{-8}$ | g/(mm<sup>3</sup>s) | [38, 41, 72] |
| Necrosis coefficient of TC | $\gamma_{\text{necrosis}}^t$ | $\sim \mathcal{U}[1, 10] \cdot 10^{-9}$ | g/(mm<sup>3</sup>s) | [38, 41, 72] |
| Oxygen consumption due to growth | $\gamma_{\text{growth}}^{nt}$ | $\sim \mathcal{U}[1, 5] \cdot 10^{-10}$ | g/(mm<sup>3</sup>s) | [38, 41, 72] |
| Intrinsic permeability of the ECM | $k$ | $\sim \mathcal{U}[0.5, 1.5] \cdot 10^{-9}$ | mm<sup>2</sup> | [41, 60, 73] |
| Poisson’s ratio of ECM | $\nu$ | $\sim \mathcal{U}[0.35, 0.48]$ | – | [38, 74, 75] |
| Shear modulus of ECM | $G$ | $\sim \mathcal{U}[120, 710]$ | Pa | [44, 60, 76–79] |
| Dynamic viscosity of ECM | $\mu^s$ | $\sim \mathcal{U}[10, 35]$ | Pa s | [44, 80, 81] |

### 2.4 Gaussian process metamodel

When a large number of model evaluations is required, the computational cost becomes prohibitive since every evaluation of the underlying tumour-growth model is expensive (an analysis needs around 30 min to simulate seven days of growth on one Intel Xeon E5-2630 v3 Haswell node (16 cores, 2.5 GHz Octocore, 64 GB RAM) in our Linux cluster). We therefore employ Gaussian processes as a metamodel (also known as a surrogate model or emulator) for approximating our porous multiphase tumour-growth model, introduced in Section 2.2.

A Gaussian process (GP) is given by

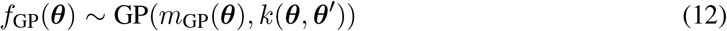

with its mean function *m*_GP_(***θ***) and its covariance function *k*(***θ***, ***θ*′**). We use a squared exponential covariance function (also called radial basis function)

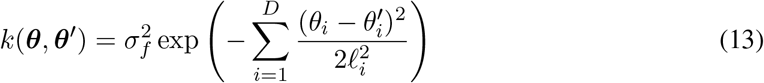

with the variance parameter *σ_f_* and one characteristic lengthscale *l_i_* per input space dimension, similar to [82]. Hence, we have *D* + 1 *a priori* unknown hyperparameters 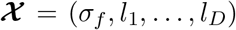. The hyperparameters 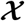 are optimised on the basis of a maximum likelihood estimation [83, 84]. To enhance the computational efficiency and robustness, we perform stochastic optimisation, using Adam optimisation [85]. Hence, training the Gaussian process means optimising the hyperparameters 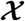.

To ensure good predictive quality of the metamodel, the training samples should provide good coverage of the input space. To this end, we use the low-discrepancy Sobol sequence [86] to generate the training samples. In addition, we generate a set 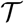 of testing samples disjoint of the training sample set. We assess the performance of the Gaussian process metamodel by evaluating the Nash–Sutcliffe efficiency *Q*^2^ [87] as

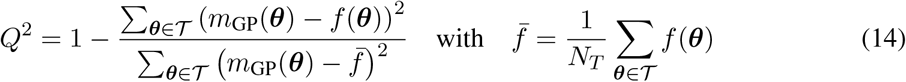

where *N_T_* is the number of testing samples. Nash–Sutcliffe efficiency compares the mean prediction of the trained Gaussian process to the actual output of the tumour-growth model, whereby values close to one indicate good agreement and thus good predictive quality of the Gaussian process.

### 2.5 Bayesian calibration

We favour a Bayesian approach to parameterise our computational tumour-growth model for a number of reasons. As stated above, the model contains uncertain input parameters for which we do not know the true value. Additionally, the true value itself may be subject to uncertainty, for instance, due to randomness in the underlying processes. This means that the uncertain input parameters are characterised by probability distributions as opposed to a single (fixed) value. When working with experimental data or biological systems, it is necessary to assume that a certain degree of uncertainty is present (such as measurement error, model error or intrinsic variability). Otherwise the solution is overconfident or incorrect. Indeed, uncertainty can also be found in computational modelling, for instance, in the form of simplifying assumptions in the model formulation. In this paper, Bayesian techniques are deemed the most appropriate for performing model calibration, since the approaches used are intrinsically able to characterise uncertainty [88].

Based on Bayes’ rule [89], knowledge of the uncertain parameters ***θ*** of a model *f* is updated in the presence of the experimental data ***y***_obs_ resulting in a posterior distribution *p*(***θ***|***y***_obs_) of the parameters as

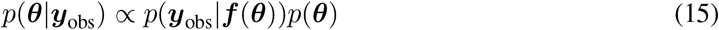

where *p*(***θ***) is the prior distribution and *p*(***y***_obs_|***θ***) is the likelihood function.

The observations ***y***_obs_ are the tumour-spheroid volumes obtained by *in vitro* experiments over time. Here, we use *N* conditionally independent observations 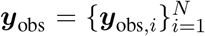, with *N* being the number of observed tumour spheroids^3^. The variable ***θ*** denotes the uncertain input parameters. We assume an additive Gaussian noise ***ϵ*** such that

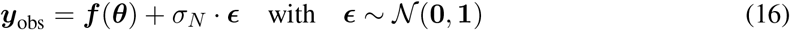

where 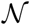 denotes the Gaussian distribution and *σ_N_* denotes the standard deviation [90].

Further, the *prior* distribution *p*(***θ***) encodes the uncertainty about the parameters before observing the data. The prior distribution is specified beforehand and represents the parameter information that we wish to include in the model calibration. It might represent the ignorance of a parameter or introduce a strong subjective belief [29]. To be consistent with the preceding GSA, we choose the prior distributions presented in Table 2. They are defined as uniform distributions, and the domains are based on values found in the literature.

The *likelihood* function *p*(***y***_obs_|***f***(***θ***)) is a true probability density for the observations ***y***_obs_, conditionally dependent on the parameters ***θ***, whereby the likelihood connects the experimental data to the computational model. In the context of Bayesian calibration, the likelihood function can be interpreted as a goodness-of-fit measure, i.e., how well the model output fits the experimental data, given a particular value for the input parameters ***θ*** [91]. Based on the additive Gaussian noise assumption (see Eq. (16)), the likelihood is given by

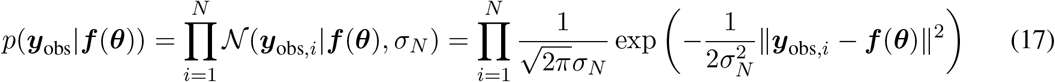

with ||·|| denoting the Euclidean L2-norm.

Finally, the *posterior* distribution *p*(***θ***|***y***_obs_) fully characterises the knowledge of the model parameters, having now observed the data. It is defined as a conditional distribution of the parameters ***θ***, given the data ***y***_obs_. The aim of this Bayesian calibration is to present the posterior distribution of the multiphase model parameters.

Due to the implicit dependency on the forward solver, the posterior distribution is analytically intractable, so it is approximated using sampling techniques [88]. However, sampling from the posterior distribution involves numerous evaluations of the forward model *f*, which results in a tremendous computational burden for complex models, such as our tumour-growth model. Therefore, the forward model *f* is again replaced by a Gaussian process *f*_GP_ (see Section 2.4) as a metamodel. This study uses a single-surface approximation [92] based on the mean of the Gaussian process. However, several approaches for capturing this additional uncertainty are proposed in [92].

We use a sequential Monte Carlo (SMC) approach [33, 93, 94] for sampling, in which the posterior distribution is approximated by a large collection of *M* ≫ 1 weighted particles [95] such that

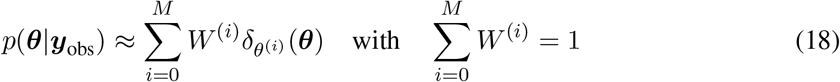

where *M* is the number of particles, 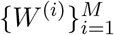 are weights, 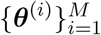 is the ensemble of particles and *δ*(·) represents the Dirac delta. The key idea of SMC is to start from a particle representation of the prior distribution and sequentially blend over to the target, i.e., posterior distribution [93]. The SMC method has gained considerable attention due to its superior efficiency and algorithmic robustness compared to Markov Chain Monte Carlo methods [93, 96, 97]. SMC comprises numerous steps, such as reweighting, resampling and MCMC-based rejuvenation^4^. The technical details of SMC are beyond the scope of this work and can be found in [33, 93, 98, 99].

### 2.6 Numerical implementation

The proposed multiphase porous media framework is implemented in the in-house parallel multiphysics research code BACI (Bavarian Advanced Computational Initiative) [100]. All methods relating to Gaussian process metamodels, sensitivity analysis and Bayesian calibration are implemented in QUEENS [101], which is a general-purpose framework for uncertainty quantification, physics-informed machine learning, Bayesian optimisation, inverse problems and simulation analytics on distributed computer systems. We use GPflow [102, 103] as the GP framework, TensorFlow [104] for Adam optimisation [85] and PyTorch [105] for generating Sobol sequences. The SMC implementation is based on the Python based SMC library *particles* [106].

## 3 Results

In this section, we present the results of the experiment, that is, the evolution of the tumour spheroid. We then move on to consider the outcome of the global sensitivity analysis and determine which model parameters are the most influential. Finally, we calibrate the relevant parameters of the multiphase model in such a way that they mimic the experimental results.

### 3.1 3D growth of neuroblastoma spheroids in microfluidic devices

To better understand how NB tumour cells interact with the microenvironment and to calibrate the tumour-growth model, we designed an *in vitro* technique based on microfluidic devices that have been described above.

The development of multicellular clusters of cells (NB spheroids) growing in microfluidic devices was evaluated by imaging to determine whether neuroblastoma primary culture (PACA cells) has the ability to form spheroids (embedded cell aggregates growing in a 3D extracellular matrix). To do this, we performed time-lapse microscopy of NB spheroids from day zero (single cells) up to day seven (Fig. 5A). The NB PACA cell culture embedded in 6 mg/mL collagen-I hydrogels induced a high degree of spheroid formation. Changes in the cell areas of NB spheroids were monitored and quantified over the culturing period (Fig. 5B): the corresponding growth curves (expressed in μm^2^) allowed us to study the size of the spheroids over time, which increased from 381±154 μm^2^ at day 0 up to 10,199±2384 μm^2^ at day 7. No significant differences were found between the biological replicates in pairing days. We closely observed the spheroid populations and determined that they displayed a narrow morphology spectrum. Specifically, a morphology analysis of individual cell clusters using bright-field imaging revealed a predominant phenotype at 6 mg/mL, with most of the spheroids presenting circular structures without elongations or protrusions on their surface. We measured the circularity of clonal cell clusters over time (Fig. 5C) and found high circularity values, i.e., high circularities of the projected area of the spheroids (with the median values ranging from 0.92 to 0.97 at various time points) confirming the predominance of a single spheroid phenotype. The lower circularity values found during the first culture days might be due to a small inherent error in measuring perimeters on small pixelated objects or the lack of total compactness during this time frame.

**Figure 5:**
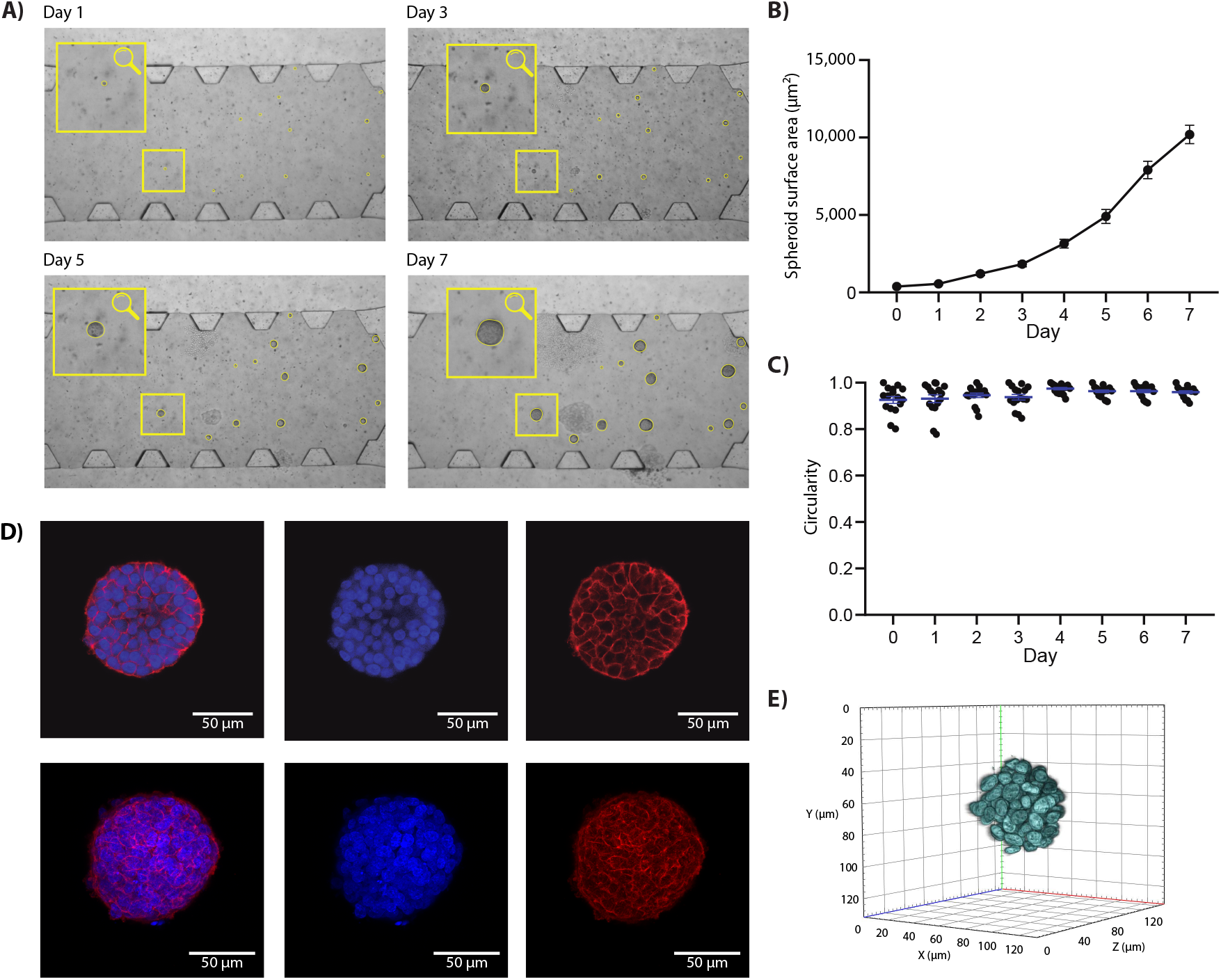
3D growth of neuroblastoma spheroids in microfluidic devices. A) Representative bright-field time-lapse images showing the formation of neuroblastoma spheroids in collagen-I on days 1, 3, 5 and 7. Yellow contours indicate the leading edges of the clusters. B) Evolution of the spheroid area of individual spheroids over time. Data is expressed as mean ± SEM (standard error of the mean). C) Evolution of the spheroid circularity of individual spheroids over time. Data is expressed as mean ± SD (standard deviation). D) Z-stack (up) and orthogonal projection (down) from NB spheroids after one week of incubation in microfluidic devices in collagen-I hydrogels. Projections were obtained from Z-stacks analysed with a Zeiss LSM 880 confocal microscope using fluorescent dyes (Merge channels (left), blue nuclear (blue nuclear counterstain with DAPI and Phalloidin-TRITC in Red to stain Actin filaments) Scale bars: 50 μm). E) 3D projection of NB spheroids containing histone 2B fused to GFP imaged with Lattice Lightsheet 7 microscope (Zeiss) lapse imaging. Images were taken on day 7, processed and 3D volume was projected with Zen 3.5 Blue software (Zeiss).

To further support our observations, we performed confocal imaging and 3D reconstructions from light-sheet microscopy. Immunofluorescence against filamentous actin (F-actin) revealed that most of the NB spheroids presented a round shape with a highly compacted cytoskeleton, very few protrusions and a smooth surface (Fig. 5D). Finally, the 3D projection displayed morphological non-invasive and rounded shape structures (Fig. 5E). This phenotype is typically associated with high cell-cell adhesion due to the high compressive forces acting on the spheroids in the 3D hydrogel. Both cell spreading and tumour invasion were greatly reduced, and most of the cells remained as dense and isolated aggregates, with minor motility through the collagen gel compared with previously studied cancer types [2].

### 3.2 Global sensitivity analysis

We now determine the first- and the total-order Sobol indices to enable us to separate uncertain input parameters of high influence from non-influential ones. Table 2 lists all eleven uncertain input parameters together with the corresponding probability distributions based on (experimental) data taken from the literature. The quantity of interest is the tumour volume.

We estimate the Sobol indices separately for each time point in the experimental measurements, i.e., from day zero to day seven. For this purpose, we train a Gaussian process in which the scalar quantity of interest is the tumour volume at the corresponding time point. This is based on 460 training samples and results in a Nash–Sutcliffe efficiency (see Eq. (14)) of the surrogate model of *Q*^2^ > 0.96 for all cases. First- and total-order Sobol indices are estimated for 1, 3, 5, and 7 days of growth.

Only five input parameters have a major impact on the tumour volume, these being the interfacial tension (*σ_tl_*), the dynamic viscosity of TC (*μ^t^*), the relative permeability exponent between the tumour cells and the ECM (*A_t_*), the tumour-growth coefficient 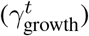 and the intrinsic permeability of the ECM (*k*). The remaining six parameters have a first- and total-order Sobol index of close to zero (see Table B1). We therefore conclude that these parameters are non-influential in the given setup and can be fixed at any value within the studied ranges.

The Sobol index estimates for the most influential parameters are shown in supplementary material Fig. 6. For all five of the most influential parameters, the total-order Sobol index is considerably higher than the first-order index. This indicates the presence of higher-order interaction effects. In this context, Sciumè et al. [38] found that changes in interfacial tension *σ_tl_* influence the speed of tumour growth. An interaction between the interfacial tension *σ_tl_* and the growth coefficient 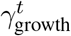 is therefore to be expected. Comparing the estimated Sobol indices for the various time points reveals that while the total-order Sobol index of the growth coefficient 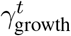 dominates the GSA, the corresponding first-order Sobol index decreases over time. At the same time, the influence of the dynamic viscosity of TC *μ^t^* increases, in particular the total-order Sobol index. This could be because the cells are initially more independent as they are separated from each other. In contrast, once the spheroid has grown, cells behave like a cluster and no longer as individual entities. Our results highlight the increasing influence of the mechanical properties of the tumour (in this case its dynamic viscosity) the longer the tumour grows. Moreover, the influence of interactions of different parameters also increases over time.

**Figure 6:**
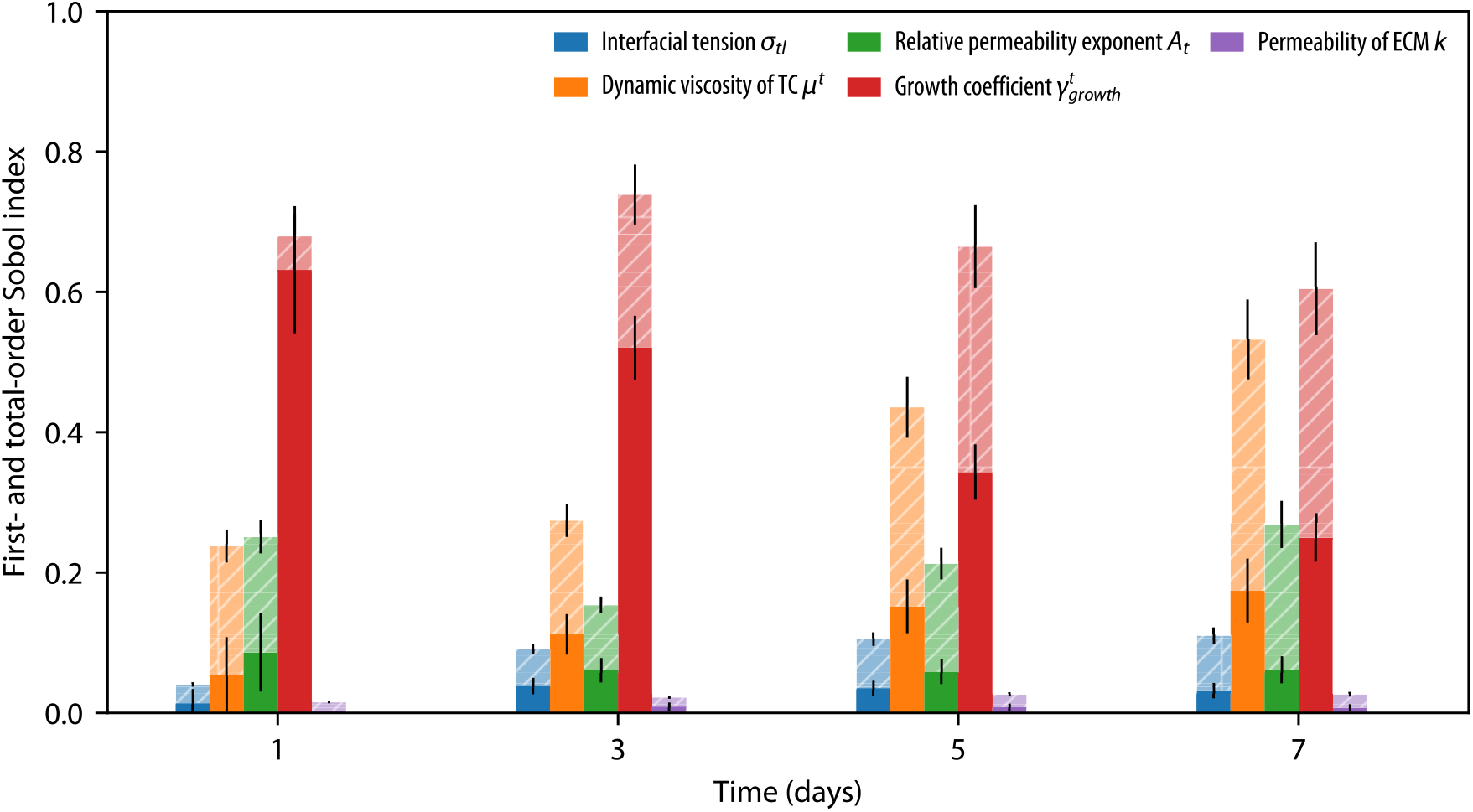
First- and total-order Sobol indices for 1, 3, 5, and 7 days of growth. First-order Sobol indices are shown as bold bars, while total-order Sobol indices are shown as dashed bars in the background. The error bars indicate 99% confidence intervals for the corresponding Sobol index estimates.

The results of the GSA show that six out of the eleven uncertain input parameters have a total-order Sobol index close to zero. This suggests that these parameters are non-influential w.r.t. the quantity of interest, i.e., the tumour volume, and can hence be fixed anywhere with the range given in Table 2. As a result, the number of input space dimensions that have to be included in the subsequent calibration can be reduced to the number of influential parameters. To validate this assumption, we fix the non-influential parameters at the mean value of the corresponding probability distributions and compare the resulting PDF of the tumour volume to the original PDF with all eleven uncertain input parameters: the resulting distributions show very good agreement for all time points (see Fig. B1), and we therefore conclude that fixing the six non-influential parameters is justified.

### 3.3 Bayesian calibration

Based on the results of the GSA, only the most influential parameters are included in the Bayesian calibration of the multiphase model. Hence, we analyse five parameters: 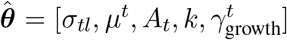. We define the priors 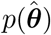 of each parameter as uniform distributions, following the ranges established in the GSA (see Table 2). The observations ***y***_obs_ are the tumour volumes measured at eight different time points in the *in vitro* spheroid experiments as described in Section 2.1. The physics-based tumour model *f* is replaced by the posterior mean of a Gaussian surrogate model *f*_GP_ trained on the following data set: the time is used as an additional input space dimension, and hence 536 training samples, each taken at eight time points, result in a training data set containing 4288 data points. To do this, we use a sparse variational Gaussian process to reduce both computational complexity and storage demands [107]. This results in a Nash–Sutcliffe efficiency of *Q*^2^ > 0.98 for the surrogate model. The posterior 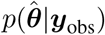 is approximated with 20,000 particles and 5 SMC rejuvenation steps. The noise variance 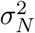 is fixed at 1×10^12^ μm^6^ (see Eq. (16)).

**Figure 7:**
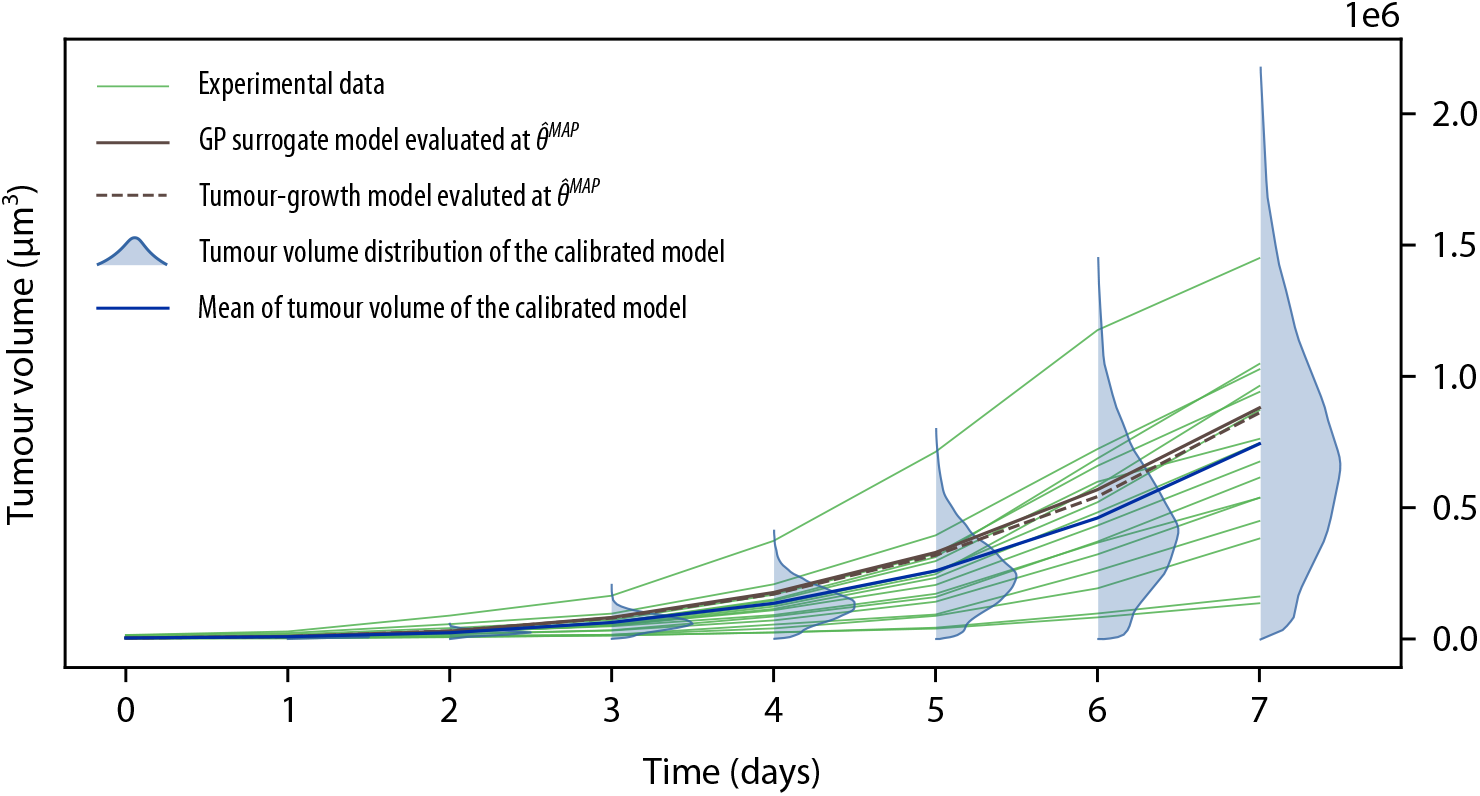
Spheroid tumour volume over time. In green solid lines, the growth of each spheroid from the experimental data set is plotted. In light green, the experimental variability is depicted. The outcome of the surrogate model evaluated at 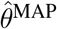 is plotted in brown, and the outcome of the forward model in dashed brown line using the same point estimate 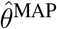. The distribution of the tumour volume *p*(*V^t^*|***y***_obs_) of the calibrated model is evaluated at each experimental time point and shown in light blue. The mean of the tumour volume of the calibrated model is plotted in dark blue.

In the following, we consider the results of the Bayesian calibration from two perspectives. First, we analyse the output distribution of the calibrated model *p*(*V^t^*|***y***_obs_) (in the output space, i.e., the tumour volume). This allows us to compare the remaining uncertainty in the tumour volume *V^t^* with the experimental data. We then analyse the posterior distribution 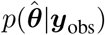 of the input parameters themselves (in the input parameter space).

The results of the Bayesian calibration in the output space are shown in Fig. 7 together with the experimental data. The combination of parameters that results in the maximum a posteriori density estimate is

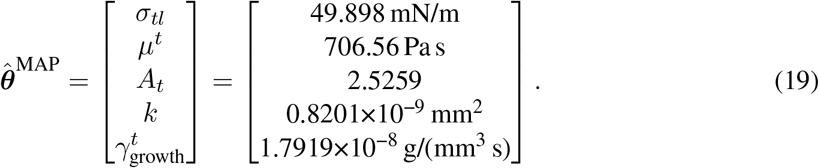

We evaluate this combination of parameters in both the tumour-growth model *f* and the Gaussian process surrogate mode *f*_GP_, and the corresponding results are depicted as brown dashed and continuous lines, respectively (Fig. 7). These plots confirm that the Gaussian process surrogate model emulates the tumour-growth model reasonably well. We also exploit the capacity of the Bayesian approach to represent uncertainty in the output space. The probability distribution of the tumour volume *p*(*V^t^*|***y***_obs_) that emerges from a forward uncertainty quantification using the posterior distribution is plotted in light blue. It estimates the probability density function of the spheroid volume over time. The mean tumour volume 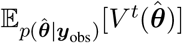 of the calibrated model is plotted in dark blue. It is noteworthy that the results of the Bayesian calibration match the experimental variability.

To gain a better understanding of the five calibrated parameters, we consider the posterior distribution 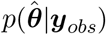 in the input parameter space. Since we are interested in the implications for each individual parameter, we first focus on 1D marginal posteriors *p*(*θ_i_*|***y***_obs_). We will then analyse the 2D marginal posteriors for parameter pairs, as the GSA indicates the presence of interactions between the input parameters. The overall results are shown in Fig. 8. It should be borne in mind that the 1D and 2D marginal distributions all are projections of the five-dimensional posterior distribution 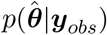 to lower dimensions.

**Figure 8:**
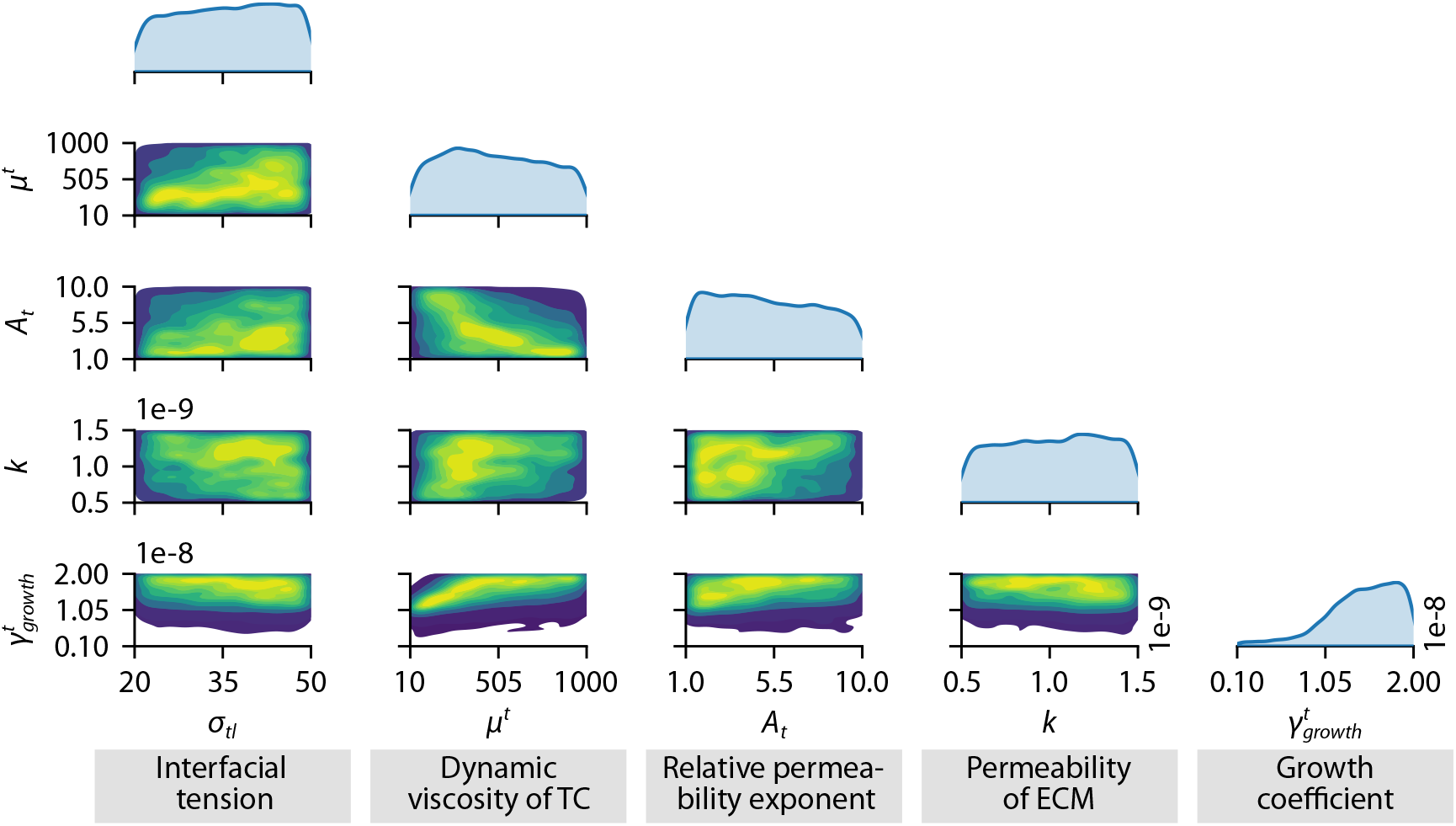
1D and 2D marginal posterior PDFs. Yellow colours indicate high values of the posterior densities, whereas purple colours indicate small values. The diagonal, in light blue, shows the 1D marginal posterior distribution for a single parameter.

The marginal posterior of the growth coefficient 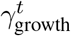 presents a higher probability mass clustered around the value 2×10^−8^ g/(mm^3^ s). Therefore, we conclude that the lower values obtained from the literature are unlikely to fit the experimental NB spheroid growth. The marginal posterior of the growth coefficient is significantly different from the other four (*σ_tl_*, *μ^t^*, *A_t_* and *k*). This is in agreement with the GSA, which indicates that the growth coefficient is the most influential parameter. Thus, we consider that the identification of a more probable range of values for this parameter is a solid outcome of the Bayesian calibration. Nevertheless, the plot does not reveal a clear peak at any certain value, presumably due to the need to explain the high variability of the experimental data.

Figure 8 further shows that the 1D marginal posterior distributions for the interfacial tension *σ_tl_* and permeability *k* of the ECM are uninformative. This finding is in agreement with the output of the GSA, in which both parameters are the least influential of the five parameters selected for the Bayesian calibration. The flat marginal posteriors indicate that no information about the effect of each individual parameter can be extracted separately from the available data. This is assumed to be due to the small influence of these parameters on the resulting tumour volume. Also, the forward model may allow for a wide range of parameters without resulting in a drastic change in the outcome. A further hypothesis is that this is due to the low data regime, which comprises only eleven spheroid data points per time step. The 1D marginal posterior distribution of the dynamic viscosity *μ^t^* reveals a slightly higher probability mass for values below 500 Pa s.

We now consider the 2D marginal posteriors. Figure 8 reveals that the dynamic viscosity *μ^t^* has a negative correlation to the relative permeability exponent *A_t_*, in other words, that large *μ^t^* values have a high density in combination with small *A_t_* values. Also, a slightly higher density occurs when high values of *σ_tl_* are combined with higher values of *μ^t^*, although low values of viscosity worked with the whole support of the interfacial tension. The 2D marginal posterior of the interfacial tension also results in higher probability values when combined with high values of *k*. Hence, future experimental and computational analysis should also consider the possible interaction of scaffold stiffness and dynamic viscosity of tumour cells. Otherwise, the combination of dynamic viscosity *μ^t^* and permeability *k* of the ECM is uninformative. The marginal posterior distribution for the interfacial tension σtl and the relative permeability exponent *A_t_* does not reveal a clearly defined peak. The 2D marginal posterior of *A_t_* and *k* shows a higher probability for low *A_t_* values.

To conclude the Bayesian calibration results, we relate them back to the theoretical framework of our computational model. In the TCAT framework, the interfacial tension (together with the interfacial curvature) causes the pressure difference between the cell phase and the surrounding medium at the microscale. However, at the macroscale, which is where all model equations are formulated, the use of Eq. (4) is only a heuristically proposed relation [35]. The interfacial tension influences the density of the tumour. Here, we only used the tumour volume for the Bayesian calibration, which is why it is not possible to distinguish between two spheroids of the same volume but with different densities. This might explain why the marginal posterior for the interfacial tension does not show a clear peak. With the inclusion of additional data, such as cell count, tumour mass or density, a more pronounced posterior may be possible. Similarly, Eq. (3) is also a heuristical relation to include interaction forces between the cell phase and the surrounding medium at the macroscale [35], which again is not consistent with the TCAT framework. Further experimental analyses of how tumour cells interact with their surroundings (ECM and other phases) are clearly necessary. From the computational side however, Miller et al. [42] have already proposed a theoretically sound version which considers the interfaces between the phases.

## 4 Summary and discussion

The current study presents a new methodology which combines *in vitro* experiments with computational modelling to investigate the growth of tumour spheroids subject to regulation by their microenvironment. We present a complete workflow, beginning with a description of the experiments, which are simulated by a multiphase poroelastic model. The model is calibrated by the experiments under uncertainty using Bayesian techniques. This multidisciplinary study integrates various technologies, which combine ideally in advancing our understanding of how *in vitro* growth of tumour spheroids is regulated by the surrounding microenvironment. The methodology we present goes one step beyond those presented in the literature [29, 30].

We start by describing the *in vitro* development and growth of tumour spheroids by means of microfluidic-based chips, with a focus on neuroblastoma cells. We employ 3D experiments, which are better able to reproduce the spatial organisation of cells and enable control of the microenvironmental conditions. We present a novel, low-cost, and accessible method for the rapid characterisation of 3D cell clusterization using microfluidic chips that allow the culture of NB cells on collagen-type I hydrogels. The potential impact of 3D cultures has strongly been emphasised previously, and authors have provided a variety of techniques for introducing 3D culture systems with the aim of attaining more reliable and comprehensive results [108]. Efforts to transfer the experiments to 3D have also been made in NB [109–113]. The hanging drop method has traditionally been used to study spheroid growth. The principle is that the cells are deposited by gravity at the bottom of the hanging droplets, where they gradually form a spheroid. However, there is no scaffold to provide support and mimic a natural tumour microenvironment for spheroids is missing, which makes this approach unsuitable for reproducing *in vivo* tumour formation. Our *in vitro* experiments were carried out in microfluidic platforms that support NB proliferation and make *in vivo* follow-up and spheroid analysis easier thanks to their good compatibility with imaging technologies. The collagen hydrogel network enhances the study of 3D growth models in a more reliable approach, and its usefulness for the study and characterisation of different tumours (including infant and brain-derived) has been widely reported [77, 114]. Previous studies have reviewed the role of ECM in NB progression, evidencing that alterations of the ECM are tumour progression mediators [79, 115]. Moreover, stiffness regulates the neuroblastoma dynamics and behaviour [77, 116] as well as the chemotherapeutic distribution and efficacy [111]. The combination of long-term 3D culture and microfluidic devices paves the way to a better understanding of tumour spheroid formation, offering a fine methodology with which to feed the computational models.

A multiphase poroelastic model based on porous media is chosen for the simulation of spheroid growth, to better account for the features of the experimental setup. A wide range of models are presented in the literature [15–18, 20]. Hydrogels are network polymeric materials with highly hydrophilic polymeric chains and are hence associated with large quantities of water, which accounts for their biocompatibility. As with most polymers, the hydrogels exhibit time-dependent mechanical behaviour due to the intrinsic viscoelasticity of the polymeric network. Therefore, we apply a viscoelastic material law for the scaffold, which seamlessly incorporates recent scientific knowledge concerning the importance of hydrogel viscoelasticity [117, 118]. To better characterise the multiphase nature of hydrated materials (like the collagen hydrogels of these spheroids), poroelastic approaches are seen as the best solution [119]. In our porohyperelastic material model, the solid phase is assumed to have a Neo-Hookean formulation (although this model could be a future line to work in) [120]. The multiphase model presented here allows us to recreate the experimental setup, as the biophysical properties of the hydrogel scaffold (i.e., stiffness, porosity and permeability) can form valuable inputs in the model. The fibre orientation of the collagen gels could form the subject of potential future research, although this is beyond the scope of this paper. Finally, studying tumour growth in a heterogeneous microenvironment (asymmetric as opposed to spherical growth) is a further area in which the experimental and computational models complement each other in a feedback loop.

Combining the experimental activity with the poroelastic model is crucial to improving our understanding of the biological processes, i.e., the onset, formation and growth of tumour spheroids. To do this, we first assess the model’s parameter sensitivity towards the relevant output quantity over time, i.e., the tumour volume. This reveals an overall dominant effect of the input parameter accounting for growth. The other influential input parameters concern the biophysical properties of the tumour cells and their interaction with the microenvironment—a fact that confirms the importance of understanding the links between cancer biophysics and biology [1]. These findings emphasise the usefulness of a genuinely global method of sensitivity analysis, such as the Sobol method, which enables detailed insights rather than local estimates. We then estimate the model parameters using Bayesian calibration, as it is intrinsically able to capture the uncertainties present in experimental measurements. Bayesian calibration also enables prior knowledge to be seamlessly integrated. The predictive probability density of the tumour volume resulting from forward uncertainty quantification using the obtained posterior clearly reflects the ability of Bayesian methodology to capture the entire *in vitro* variability. In future studies, the Bayesian approach offers a natural way of integrating additional data as it becomes available: knowledge of the uncertain input parameters can be further updated using the posterior obtained in the present study as a prior. Further research could entail modelling other sources of uncertainty, such as the effect of spatial- and/or time-variable nutrient distribution. Such effects can be included in the analysis as additional random variables, in a similar way to [121], and might indeed result in a more expressive posterior.

This study aims at gaining a better understanding of how spheroid growth evolves over time and how it is regulated by mechanical stimuli. In this context, we are able to monitor tumour-spheroid growth under physiological conditions using 3D microfluidic devices and collagen hydrogels to mimic the ECM. Nevertheless, measuring certain properties of cells or organoids is unfeasible in such experiments. To characterise the mechanical properties of the spheroids, they usually must be separated from the collagen network and subjected to further tests, such as atomic force microscopy. This is rather complicated and no longer physiological in nature, because cells have to be separated from the organoid. The presented workflow may be more appropriate, as it not only allows the properties of the whole spheroid to be estimated but also focuses on the properties of the cells embedded in the spheroid.

We have thus shown that we are indeed able to characterise the NB spheroids’ temporal evolution, which additionally allows us to indirectly characterise their mechanical properties. In this study, we were able to estimate the permeability of both ECM and tumour cells, as well as the viscosity of the latter and the interfacial tension between the cells and the IF. The workflow described here hence enables an indirect measurement of the mechanical properties of tumour-spheroid growth purely from the images acquired directly in the course of the experiment. Additionally, being able to replicate the experimental outcomes leads to a significant reduction in cost, since it allows new experimental scenarios to be tested computationally *a priori*. This is of major importance when we study infant cancer, where early diagnosis is key. In paediatric malignancies, trial sample sizes are kept as low as possible while maintaining the ability and power to address the scientific objectives of interest. Therefore, the availability of childhood cancer cells, and here specifically NB cells, is low. This also leads us to the conclusion that computational modelling has an important role to play in the exploration of these cancers.

## 5 Acknowledgements

SHR, PG, JFMS, MJGB and JMGA were supported by PRIMAGE (PRedictive In-silico Multiscale Analytics to support cancer personalized diaGnosis and prognosis, empowered by imaging biomarkers), a Horizon 2020—RIA project (Topic SC1-DTH-07-2018), grant agreement No. 826494. SHR gratefully acknowledges the support of the Government of Aragon (Grant no 2019-23), the Deutscher Akademischer Austauschdienst (DAAD-91819992), the Fundación Ibercaja-Cai (No IT 5/21, IT 1/22) and the “Iberus+” project, co-funded by the European Union’s Erasmus+ programme and managed by Campus Iberus. Authors would like to acknowledge the use of Servicio General de Apoyo a la Investigación-SAI (Universidad de Zaragoza) and the use of Servicios Científico Técnicos del CIBA (IACS-Universidad de Zaragoza). BAS was supported by the Visiting Fellowship for Alumni Fellows of the Institute for Advanced Study (IAS). GRR was supported by the German Federal Ministry of Education and Research (project FestBatt 2, 03XP0435B). JN and WAW wish to acknowledge funding of the Deutsche Forschungsgemeinschaft (DFG, German Research Foundation) via project WA 1521/23. WAW was supported by BREATHE, a Horizon 2020—ERC–2020–ADG project, grant agreement No. 101021526-BREATHE. This work is part of a project that has received funding from the European Research Council (ERC) under the European Union’s Horizon 2020 research and innovation programme (ICoMICS grant agreement No 101018587)”.

## A Porous multiphase tumour-growth model equations

The main equations of the tumour-growth model will now be presented.

### ECM

The ECM as the solid phase is governed by the balance of momentum

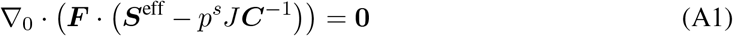

with the right Cauchy-Green tensor ***C*** and the Jacobian determinant *J* of the deformation gradient ***F***. Further, ***S***^eff^ denotes the effective second Piola-Kirchhoff stress tensor, and the solid pressure contribution *p^s^* = *S^t^p^t^* + *S^l^p^l^* (where *S^t^* and *S^l^* are the tumour cells and interstitial fluid phase saturation, respectively) [41].

We assume a visco-elastic constitutive law for the solid phase. Its hyperelastic part is assumed to follow a Neo-Hookean model with a strain energy function given by

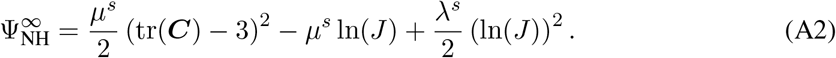

The Lamé constants in the Neo-Hookean model are defined as

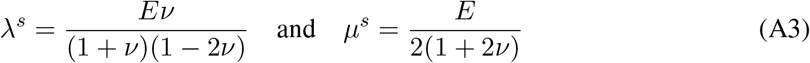

with the initial Young’s modulus *E* and Poisson’s ratio *v*. The viscous part is added to the hyperelastic part in the form of a generalised Maxwell model [46], and we assume that a dashpot in parallel with a spring is described by the following strain energy function

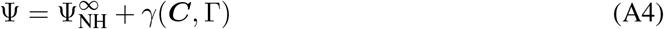

with the dissipative potential *γ*. This results in the fictitious stress tensor ***Q*** and its kinematic conjugate Γ

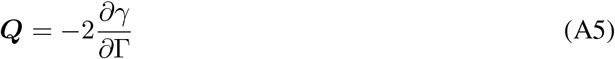

similar to the right Cauchy-Green strain tensor ***C*** and its kinematic conjugate being the second Piola-Kirchhoff stress tensor ***S***. The Maxwell model follows as

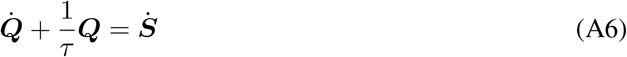

with the relaxation time *τ*. The global stress tensor is thus given by ***S*** = ***S***^∞^ + ***Q***.

Based on the shear modulus *G* and the dynamic viscosity of the ECM *μ^s^*, we calculate

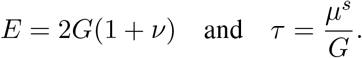

### Fluid phases

The set of primary variables of the fluid phases is given as ***ψ*** = [*p^tl^, p^l^*] with the differential pressure between the two fluid phases *p^tl^* and the pressure of the medium *p^l^*. The mass balance equation of tumour cells is then given by

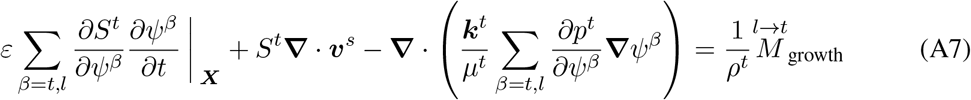

and of the medium

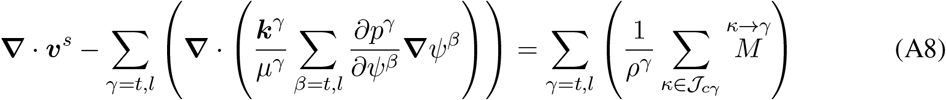

with the mass transfer terms on the right-hand side of the equation summarised in Table A1.

**Table A1:**
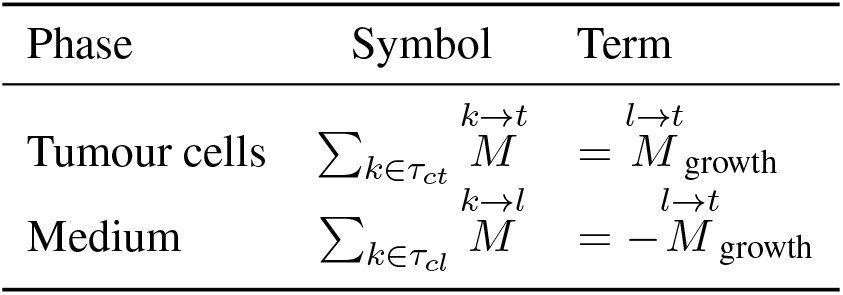
Mass transfer for the different phases of the tumour-growth model.

### Species

We consider oxygen (denoted by the superscript *n*) to be the only nutrient which governs the growth of the tumour. Oxygen is transported in the medium and limits the growth of the tumour spheroid. The mass balance equation for oxygen with mass fraction 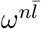 is given by

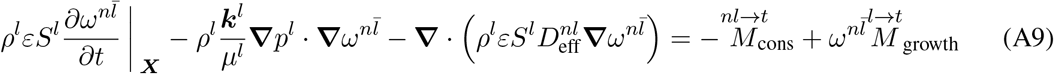

where the terms on the right-hand side of the equation describe the mass transfer of species between the different phases.

We further include necrotic tumour cells (denoted by the superscript *N*) as part of the tumour cell phase. Assuming that the cells cannot diffuse, the mass balance equation for necrotic tumour cells reduces to

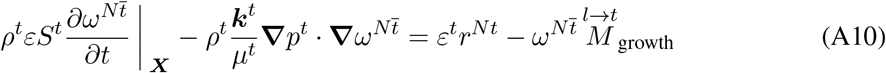

### Computation of tumour volume

The tumour volume is given by

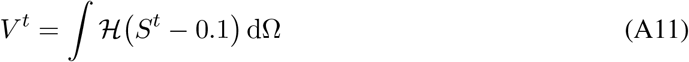

with the Heaviside function 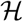. The tumour is defined as the part of the domain where the saturation of tumour cells is greater than 0.1 (*S^t^* > 0.1).

## B Additional results

**Table B1:** First- and total-order Sobol indices. All values are rounded to three decimal places.

| Parameter | Day 1 |  | Day 3 |  | Day 5 |  | Day 7 |  |
| --- | --- | --- | --- | --- | --- | --- | --- | --- |
| | $\mathcal{S}_i$ | $\mathcal{S}_i^{tot}$ | $\mathcal{S}_i$ | $\mathcal{S}_i^{tot}$ | $\mathcal{S}_i$ | $\mathcal{S}_i^{tot}$ | $\mathcal{S}_i$ | $\mathcal{S}_i^{tot}$ |
| $\sigma_{tl}$ | 0.014 | 0.040 | 0.038 | 0.091 | 0.035 | 0.105 | 0.032 | 0.110 |
| $\mu^t$ | 0.055 | 0.238 | 0.112 | 0.274 | 0.152 | 0.436 | 0.175 | 0.532 |
| $A_t$ | 0.087 | 0.251 | 0.061 | 0.154 | 0.059 | 0.213 | 0.062 | 0.269 |
| $A_l$ | 0.0 | 0.0 | 0.0 | 0.0 | 0.0 | 0.0 | 0.0 | 0.0 |
| $\gamma_{growth}^t$ | 0.632 | 0.679 | 0.521 | 0.739 | 0.344 | 0.665 | 0.250 | 0.605 |
| $\gamma_{necrosis}^t$ | 0.0 | 0.0 | 0.0 | 0.0 | 0.001 | 0.001 | 0.001 | 0.001 |
| $\gamma_{growth}^{nt}$ | 0.0 | 0.0 | 0.0 | 0.001 | 0.0 | 0.002 | 0.0 | 0.001 |
| $k$ | 0.003 | 0.015 | 0.009 | 0.023 | 0.008 | 0.027 | 0.007 | 0.027 |
| $\nu$ | 0.001 | 0.003 | 0.0 | 0.0 | 0.0 | 0.001 | 0.0 | 0.001 |
| $G$ | 0.001 | 0.004 | 0.0 | 0.001 | 0.0 | 0.004 | 0.0 | 0.005 |
| $\mu^s$ | 0.0 | 0.0 | 0.0 | 0.0 | 0.0 | 0.0 | 0.0 | 0.0 |

**Figure B1:**
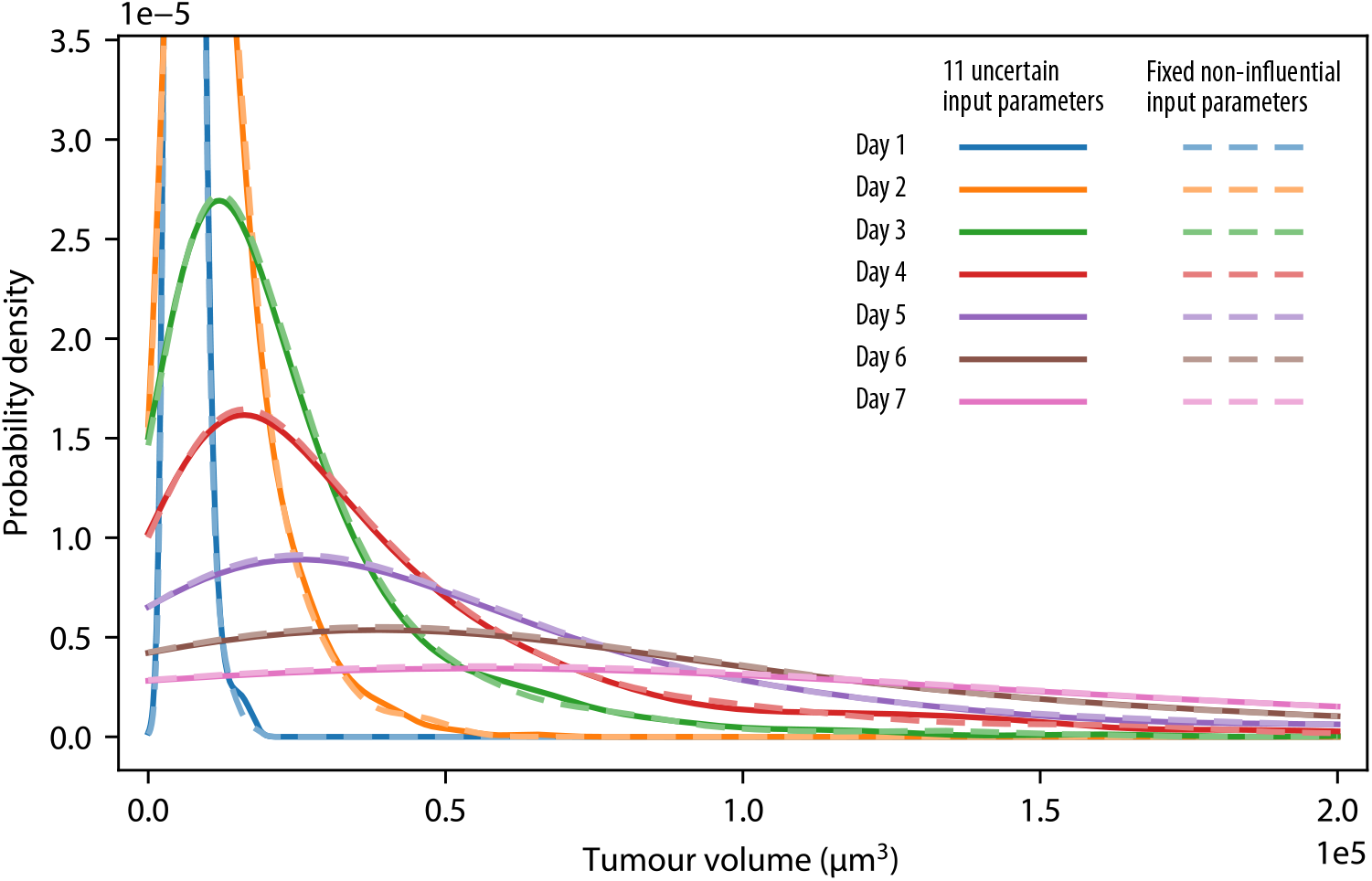
Probability density functions (PDFs) for different numbers of uncertain input parameters. The solid lines show the distributions with all 11 uncertain input parameters as listed in Table 2. The dashed lines represent the distributions with the six non-influential parameters fixed at their mean values.

**Table B2:**
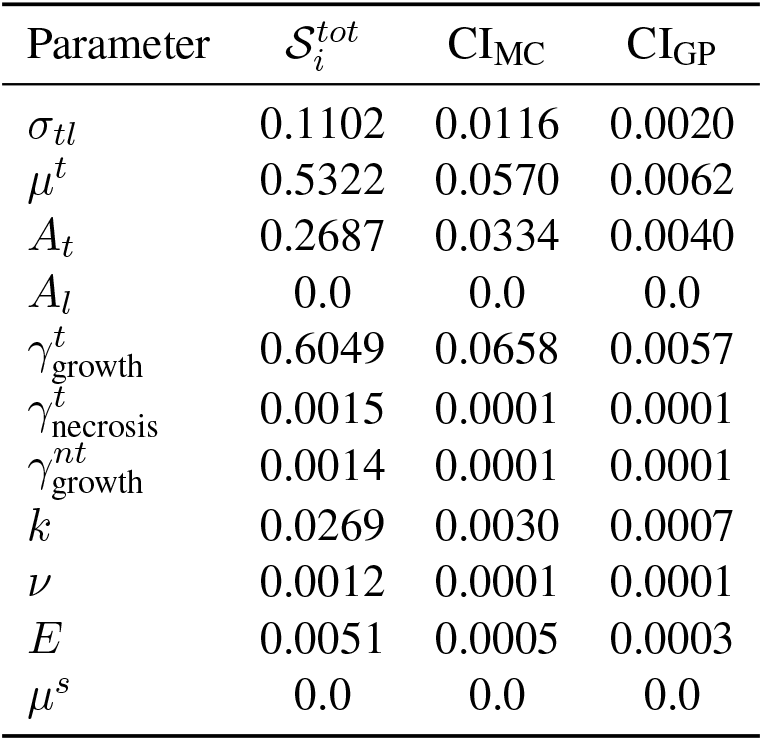
99% confidence intervals (CI) due to Monte Carlo (MC) integration and the use of the Gaussian process (GP) metamodel for total-order Sobol index at day 7 rounded to four decimal places. Estimation based on 10,000 Monte Carlo samples, 300 bootstrap samples, and 500 realisations of the trained Gaussian process surrogate.

## Footnotes

1 In [21, 23, 24, 35–41], the superscript *l* denotes the interstitial fluid which surround cells in the body and transports nutrients. The IF directly corresponds to the culture medium in the context of *in vitro* spheroid growth, and we therefore use the same superscript *l*.

2 Hereafter, we will refer to the Sobol indices as 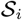. Although the notation is similar to that for saturation (*S^α^*) of the fluid phases of the porous media model, it was nevertheless decided to retain it since both are standard notations in their corresponding fields.

3 To avoid cluttered notation, the output of ***f***(***θ***) is a vector containing the tumour-spheroid volumes at different discrete time points. Similarly, ***y***_obs, *i*_ contains the experimentally observed volume of one tumour spheroid *i* at different discrete time points, and ***y***_obs_ concatenates the tumour-spheroid volumes for all time points and all observed spheroids.

4 In the rejuvenation step, we use a random walk Metropolis kernel as the MCMC transition kernel, where the covariance matrix of the proposal distribution is set proportionally to the empirical variance of the particles [94]. An adaptive tempering scheme [94] was employed such that the ratio between two subsequent tempering exponents equals 0.95.

## Notes

### Competing Interest Statement

The authors have declared no competing interest.

